# MVPA does not reveal neural representations of hierarchical linguistic structure in MEG

**DOI:** 10.1101/2021.02.19.431945

**Authors:** Sophie Arana, Jan-Mathijs Schoffelen, Tom Mitchell, Peter Hagoort

## Abstract

During comprehension, the meaning extracted from serial language input can be described by hierarchical phrase structure. Whether our brains explicitly encode hierarchical structure during processing is, however, debated. In this study we recorded Magnetoencephalography (MEG) during reading of structurally ambiguous sentences to probe neural activity for representations of underlying phrase structure. 10 human subjects were presented with simple sentences, each containing a prepositional phrase that was ambiguous with respect to its attachment site. Disambiguation was possible based on semantic information. We applied multivariate pattern analyses (MVPA) to the MEG data using linear classifiers as well as representational similarity analysis to probe various effects of phrase structure building on the neural signal. Using MVPA techniques we successfully decoded both syntactic (part-of-speech) as well as semantic information from the brain signal. Importantly, however, we did not find any patterns in the neural signal that differentiate between different hierarchical structures. Nor did we find neural traces of syntactic or semantic reactivation following disambiguating sentence material. These null findings suggest that subjects may not have processed the sentences with respect to their underlying phrase structure. We discuss methodological limits of our analysis as well as cognitive theories of “shallow processing”, i.e. in how far rich semantic information can prevent thorough syntactic analysis during processing.

## 1. Introduction

Although we perceive language mainly in a sequential fashion (e.g. by reading word by word) we need to take into account information beyond the sequential order to fully comprehend its meaning. For example, in a sentence like “The woman who owns two dogs chases the cat” we understand that the woman is the one chasing, not the dog. This knowledge can be expressed through hierarchical, structured relationships between the words. Specifically, words can be grouped into constituents (e.g. “Who owns the dog” and “The woman chases the cat”) and constituents in turn can be nested into higher-level phrases, as shown in 1. The resulting nested phrase structure then fully describes the important conceptual units and their relationships with each other. Thus, hierarchical phrase structure also directly relates to thematic role assignment (the woman being assigned the agent role of the chasing action).

1. ((The woman (who owns the dogs)) chases the cat)

This type of structured meaning is to a large degree determined by syntax. As seen above, syntactic aspects like word order, function words (here: the relative pronoun ‘who’) as well as morpho-syntactic features such as number agreement provide cues with respect to the word-phrase relationships. Semantic information (e.g. animacy) or even just semantic association itself can also guide how structure should be assigned. In the above example, syntactic cues, however, override simple semantic association between the lemmas “dog” and “chase”. In theory, hierarchical descriptions can be applied to all linguistic levels of the stimulus during language processing (e.g. syntactic, semantic and phonological structure) (Jackendoff [2003]).

How hierarchical phrase structure building is neurally encoded as we process language is still an open question. In fact, some have even disputed its neural and psychological reality during language use altogether (Frank et al. [2012]). Some recent evidence for the reality of hierarchical phrase structure building comes from neuroimaging studies that assess its consequences on memory load (Nelson et al. [2017]; Pallier et al. [2011]) and production (Giglio et al. (in prep)). For example, Pallier et al. varied linguistic constituent size while keeping overall sentence length constant and identified brain regions whose activity parametrically increased with the size of the constituents (larger constituents thought to result in higher memory demands and stronger neural activity) (Pallier et al. [2011]). Following a similar approach, Nelson et al. modelled neural activity according to a hierarchical phrase-structure model and found it to explain more variance when fitted to intracranial data as compared to alternative models that were based on transition probabilities only (Nelson et al. [2017]). This is in line with behavioral evidence, demonstrating that humans prefer a hierarchical interpretation over a linear one, for example when interpreting ambiguous noun phrases, such as “second blue ball” (Coopmans et al. [2021]). At the same time, there are several studies demonstrating that reading times can often be sufficiently accounted for by sequential-structure models (Frank and Bod [2011]), casting doubt on how pervasive the construction of hierarchical structure during language processing really is.

In early psycholinguistic experiments, hierarchical structure building has been measured through reading time behaviour for structurally ambiguous sentences. One example for such ambiguity is prepositional phrase attachment. Prepositional phrases (PPs) in sentence-final position (examples 2 & 3) are structurally ambiguous with respect to their attachment to the main clause. For example, a prepositional phrase can be interpreted as noun-attached as in sentence 2 (a cop with the revolver) or as verb-attached as in sentence 3, in which case it modifies the verb (seeing with binoculars). In contrast to other structurally ambiguous stimuli such as garden-path sentences, different prepositional phrase attachments do not involve different word forms or function words. Hence, any disambiguation cannot depend on syntactic information. Still, human readers are able to assign a unique meaning to such structurally ambiguous sentences with ease, relying on world knowledge to connect the semantic information provided by both the prepositional phrase itself with its preceding context in the most plausible way (e.g. revolvers are likely to be carried by cops and binoculars are likely instruments for seeing.). Note that sentence-final prepositional phrases are not rare or non-canonical. For example, in the structurally annotated TIGER corpus (see methods for details) we found about 43% of all prepositional phrases to be structurally ambiguous.

2. The spy saw the cop with the revolver.

3. The spy saw the cop with the binoculars.

Originally, structurally ambiguous sentences had been shown to lead to prolonged reading times at the disambiguating word (e.g. noun-attached PPs being read more slowly than verb-attached PPs). Based on these findings, Frazier had proposed sentence comprehension to rely on an initial structural interpretation of the sentence driven by syntactic cues only and following certain rules such as the minimal attachment principle. According to the minimal attachment principle, the preferred structure is always the more shallow one (i.e. the one resulting in a minimal amount of nested dependencies). Therefore, according to minimal attachment the verb-attached reading of the PP is preferred already when encountering the preposition. In the case of a noun-attached phrase, subsequent words thus leads to the need for post-hoc structural reanalysis and as a consequence longer reading times (Rayner et al. [1983]; Frazier and Rayner [1982]). Frazier’s early theory was quickly overturned in favour of a parallel (or cascading) processing model(McClelland and Kawamoto [1986]; Van Den Brink and Hagoort [2004]; Pulvermüller et al. [2009]; Hagoort [2017]) by several studies demonstrating the fast integration of non-syntactic cues early during online processing (Spivey-Knowlton and Sedivy [1995]),(Altmann and Steedman [1988]),(Taraban and McClelland [1988]),(Traxler and Tooley [2007]),(Mohamed and Clifton [2011]). For the processing of ambiguous PPs, it has been shown that facilitated processing of verb-attachments is modulated by referential information imposed by the context (Altmann and Steedman [1988]) as well as semantic content of the preceding verb (Spivey-Knowlton and Sedivy [1995]). More concretely, Spivey-Knowlton et al. have shown that action verbs bias expectations towards verb-attachment while verbs referring to mental states (e.g. the spy hoped for ..) or perception can bias towards noun-attachment (Spivey-Knowlton and Sedivy [1995]). The authors explain this by different types of verbs being associated with certain thematic roles to different degrees (e.g. action verbs occur with an instrument more often than perception verbs). As a consequence, reading time differences that have originally been interpreted to be a direct consequence of hierarchical structure building, could be reflecting predictions about upcoming semantic content instead.

In a more recent study, Boudewyn and colleagues argued against this alternative hypothesis of PP reading differences being caused by varying semantic predictions. They investigated the neural activity evoked by verb- and noun-attached prepositional phrases through event-related potentials (ERPs). In addition to the classically observed delay in reading times, their noun-attached stimuli evoked larger positive potentials around 600 ms (P600) (as compared to their verb-attached versions). Importantly, they showed that the amplitude of this P600 was reduced when noun-attached targets followed noun-attached primes (Boudewyn et al. [2014]). Boudewyn and colleagues are not the first ones to report structural priming effects. In fact, syntactic priming has been reported already some 35 years ago, showing that speakers are more likely to repeat a given syntactic structure in their utterances than to switch between two conceptually equal alternatives (Bock [1986]). To evoke priming of hierarchical structure, researchers explicitly vary lexical information while keeping syntactic structure stable. More recent investigations indicate, however, that event structure (i.e. thematic roles) as well as lexical information can to a large degree account for many priming results and hence priming solely on the structural level has not been definitively proven yet (Ziegler et al. [2019]). Other confounding factors that can evoke priming and are often contrasted along side syntactic structures are information structure, syntax-animacy mapping and rhythmic priming. Boudewyn et al. argue for their priming effect to be structural in nature based on the timing of their observed ERP effect. Differences in ERPs have been generally interpreted as neural markers for a difference in processing (for example more or less engagement of the underlying neuronal population). The P600, specifically, has been reported most often in the context of syntactic violations or anomalies. Hence, the authors interpret this priming effect to reflect facilitated structural processing of an originally dis-preferred structure. Still, ERP effects need to be interpreted with caution, since their relationship to underlying cognitive mechanisms is unclear. For example, recent computational cognitive models of language processing illustrate that ERP markers can be modelled as reflecting general update or error signals, without restricting them to any specific linguistic operation (Rabovsky et al. [2018]),(Fitz and Chang [2018]).

In addition, most ERP research so far reflects only a one-sided measure of the neural code. Namely, the dominant analysis approach has been to treat ERPs as unidimensional point-estimates. Computing signal amplitude separately for a given channel and time point and averaged over trials, subjects and eventually space and time. As a consequence, such analyses can only detect univariate effects and are highly sensitive to subject-level variability. With the recent increase in computing power and developments of multi-variate pattern analysis (MVPA) we can now capture richer multidimensional information encoded across several channels or source points (Guggenmos et al. [2018]; Norman et al. [2006]). Through MVPA, researchers have been able to uncover additional task-relevant brain regions (Jimura and Poldrack [2012]) and characterise the specific computations needed for ambiguity resolution in more detail (Tyler et al. [2013]). Furthermore, MVPA has the potential to be sensitive to distributed neural representations of the content whereas univariate methods have been thought to be most sensitive to the engagement of basic processing operations (Raizada et al. [2010], Mur et al. [2009], Okada et al. [2010]). Although not every effect revealed through MVPA is necessarily indicative of an underlying distributed neural code (Davis et al. [2014]), the technique has nonetheless been successfully used to reveal higher-level structure in the neural signal for domains other than language (e.g. for hierarchical motor sequences Yokoi and Diedrichsen [2019]). MVPA might hence be better suited to target hierarchical structure building during language processing than previous univariate methods.

In this study, we revisit processing of structurally ambiguous PPs with the approach of MVPA in order to more directly tap into representations of hierarchical structure underlying language comprehension. In contrast to early psycholinguistic approaches we do not assume that noun or verb-attached prepositional phrases are processed differently from each other in the sense of one structure being more preferred over another. Rather we ask, whether it is possible to find a neural correlate of the hierarchical phrase structure of a sentence (i.e. neural patterns that distinguish between verb- and noun-attached PPs), given completely ambiguous syntactic cues.

## 2. Methods

### 2.1. Stimulus Material

#### 2.1.1. Corpus Analysis

All stimuli were created in German. Since most of the previous literature had looked at prepositional phrases in English, we first conducted a corpus analysis to determine which German preposition will most likely be ambiguous with respect to structural attachment of the prepositional phrase. For our corpus analysis we used the TIGER corpus, a manually annotated corpus of 40,000 German sentences (Brants et al. [2004]). The corpus is available at www.ims.uni-stuttgart.de in both xml as well as conll09 format. We used the xml version for queries with the TIGERSearch Tool as well as the conll09 version for quick extraction of frequency statistics using the bash shell command awk. We extracted separate frequency information per preposition and structure (noun-attached and verb-attached prepositional phrases) through the TIGERSearch software (see Appendix for details on the TIGERSearch queries).

#### 2.1.2. Stimuli

Based on the corpus search, we selected the preposition “mit” (engl.: with) because it occurs with high frequency (Figure 1) and equally often within both noun- and verb attached phrases (Figure 2). We created a stimulus set of 100 sentence pairs in German. All sentences consisted of nine words each, a subject-verb-object structure in the main clause followed by a four word prepositional phrase including the preposition and a determiner- adjective-noun phrase. This sentence structure was syntactically ambiguous with respect to the attachment site of the prepositional phrase. Within a given pair, the same prepositional phrase was presented while the sentence context leading up to it was manipulated. Based on the combined semantic information of the sentence context and the prepositional phrase, the interpretation of the most plausible attachment could be disambiguated. To steer the preferred attachment interpretation, we manipulated the sentence context in two ways. In half of the sentence pairs we varied the main verb, we call this the verb condition (examples 4 & 5). Sentence pairs in the verb condition were constructed such that the noun in object position could potentially be modified by the PP but did not have a particularly strong semantic association with the PP internal noun. By presenting these sentences with a verb for which modification through the PP internal noun was either allowed or forbidden (or at least unlikely), a verb-attached interpretation could either be encouraged or prevented respectively. In the other half of the sentence pairs, we exchanged agent and patient identity across the two sentences. In the following, I will refer to this as the role condition (examples 6 & 7). For sentence pairs in the role condition the two nouns preceding the PP had a varying degree of semantic association to the PP internal noun while the verb was held stable with a mild semantic association to the PP internal noun and optional modification through a PP. This lead to a noun-attached interpretation if the more strongly associated noun occurred in object position (the noun immediately preceding the PP) but to a verb-attached interpretation when it occurred in subject position. In both the role and the verb condition, each verb was repeated exactly two times across all sentences. We explore difference between verb and role conditions in the behavioral data but collapse across both conditions when analysing the neural data. Finally 100 filler sentences with varying syntactic structure were created.

**Figure 1:**
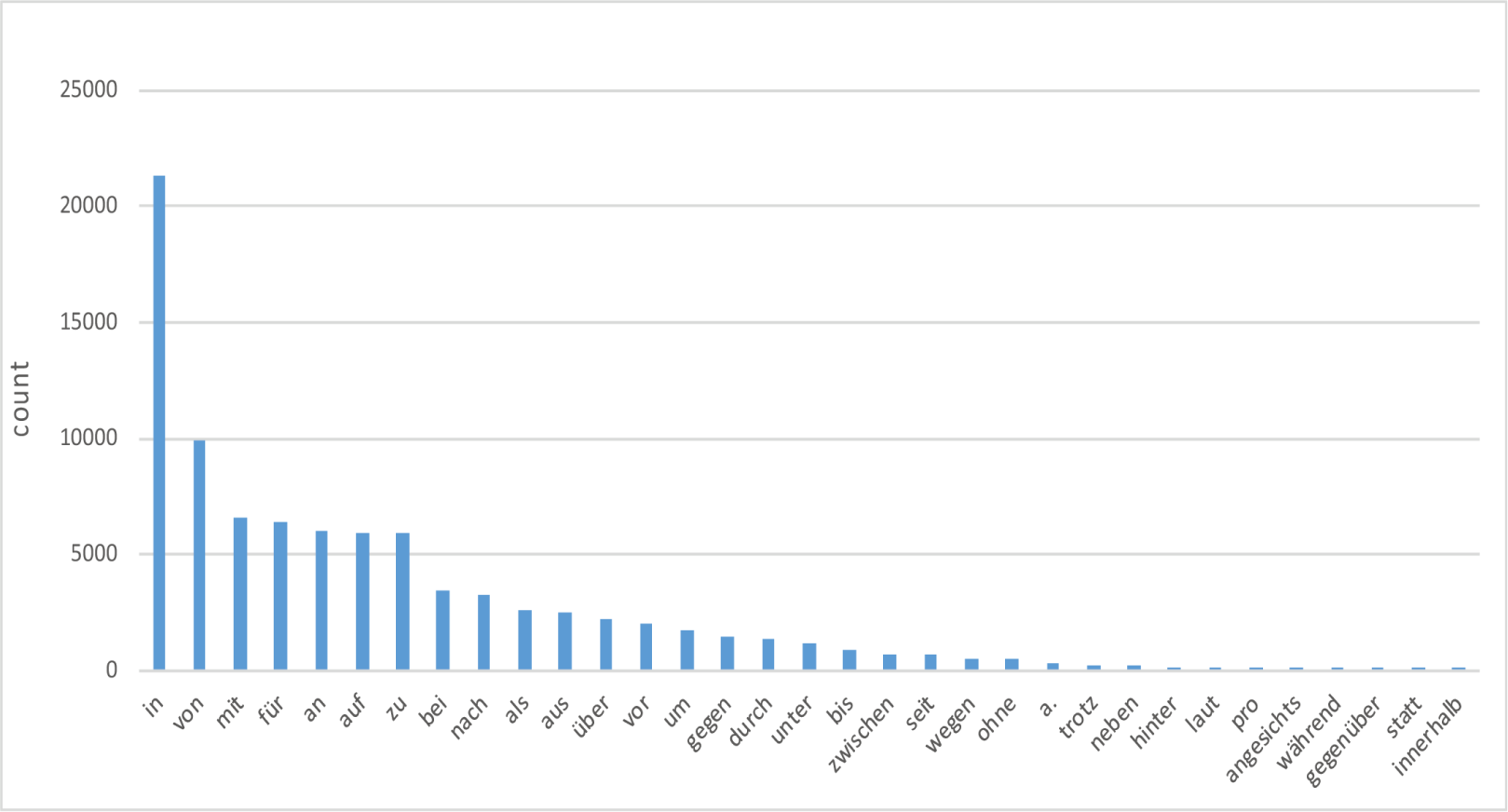
Tiger corpus frequencies per preposition. Total number of occurrence for the 33 most frequent prepositions based on the German “Tiger” corpus.

**Figure 2:**
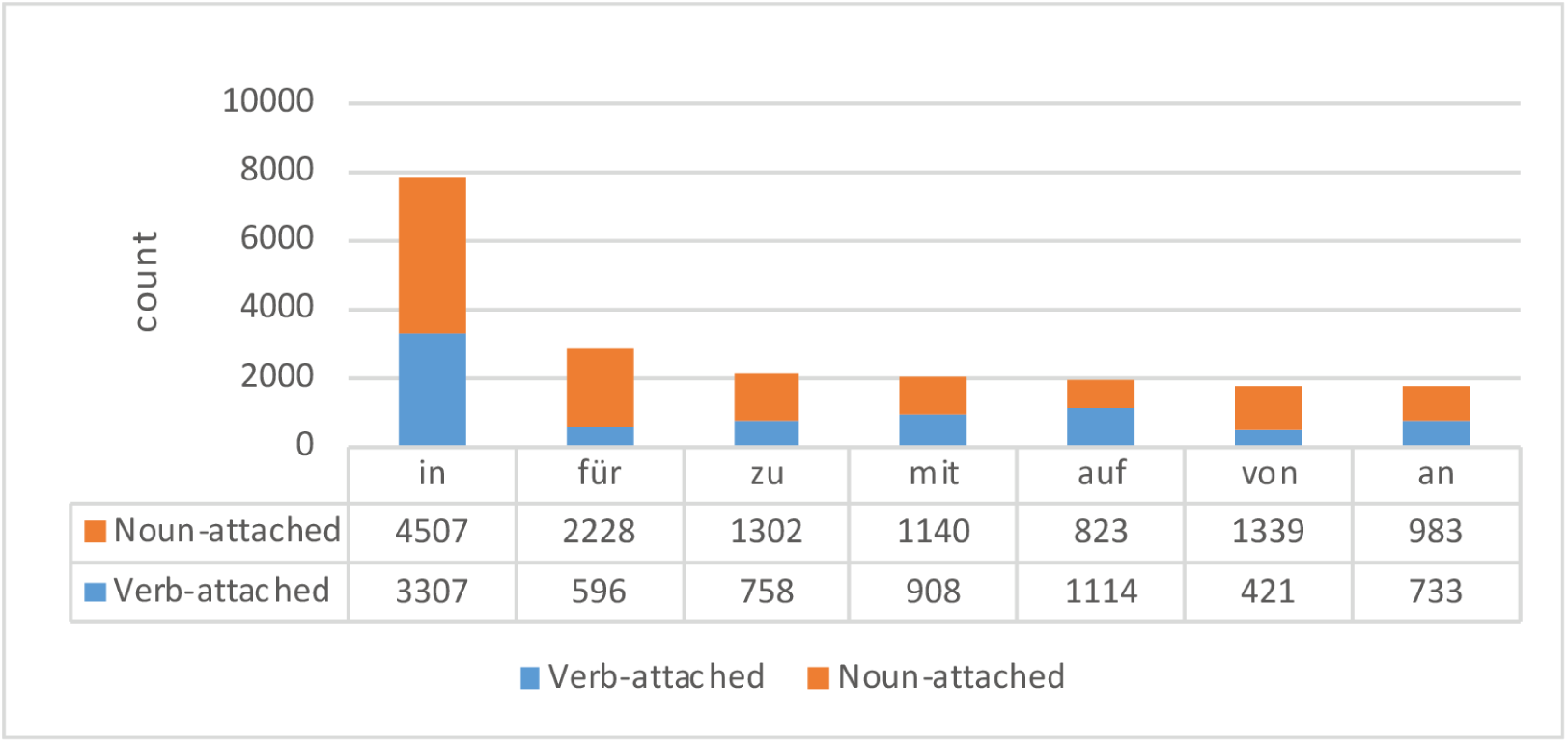
Tiger corpus attachment proportions per preposition. Frequency of verb- and noun-attached phrase constructions (not restricted to sentence final PP) for the seven most frequent prepositions in the corpus.

##### Verb condition

4. Die Partei besitzt eine Untergruppe mit einigen Argumenten

*engl.: The party has a subgroup with questionable arguments*.

5. Die Partei überzeugt eine Untergruppe mit einigen Argumenten

*engl.: The party convinces a subgroup with questionable arguments*.

##### Role condition

6. Das Kind verängstigt das Insekt mit dem giftigen Stachel

*engl: the child frightens the insect with the poisonous sting*

7. Das Insekt verängstigt das Kind mit dem giftigen Stachel

*engl: the insect frightens the child with the poisonous sting*

#### 2.1.3. Pre-test

For the majority of the sentences, the overall semantics licensed both PP attachments, even if they were constructed such that one attachment should be perceived as more plausible. To verify that our manipulation evoked the intended sentence interpretation we pre-tested all stimuli via an online questionnaire, created with the survey tool Limesurvey (Carsten Schmitz [2012]). During this online questionnaire, 62 native German speakers with a mean age of 25 (range 19-33) judged for each stimulus-sentence whether it contained a verb- or noun-attached prepositional phrase and how plausible they found the sentence (on a scale from 1 to 5). All subjects gave informed consent prior to filling in the survey and received financial reimbursement. Based on the answers we selected 200 sentences out of a larger set of 469 sentences according to criteria described in detail below (see Table 2 and 3 in Appendix for the final selection of sentences as used in the MEG experiment).

**Table 1:**
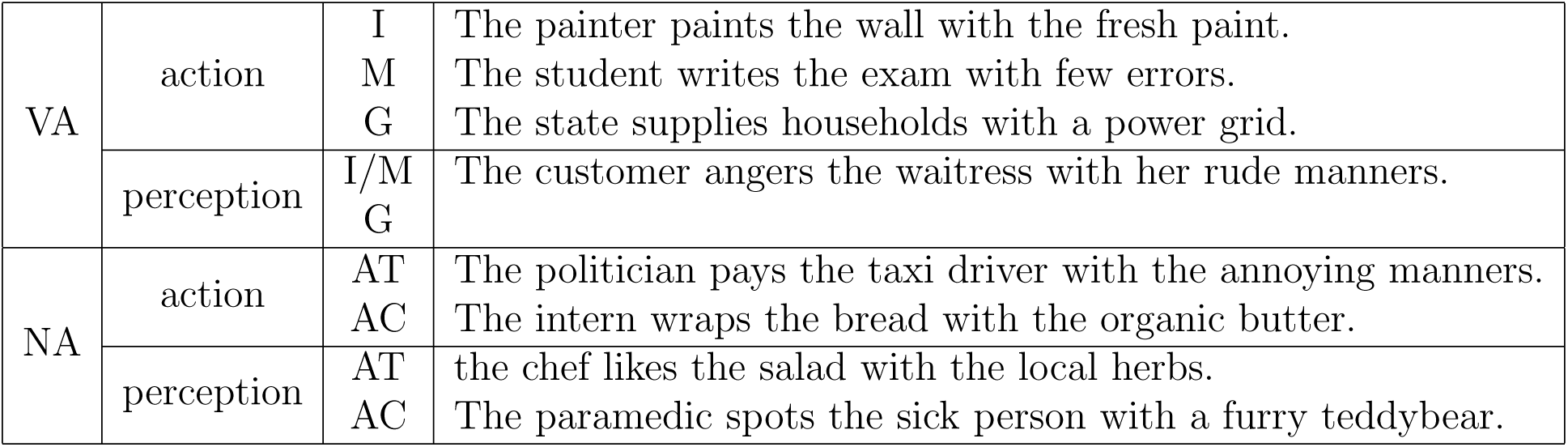
Example sentences. For each verb-attached (VA) or noun-attached (NA) PP several thematic roles could occur within the stimuli. Possible roles are instrument role (I), manner role (M), goal role (G), attribute role (AT) accompanying role (AC). Categorisation of thematic roles following those in Taraban and McClelland [1988]

First, subjects were instructed about the difference in attachments. This was done using unambiguous stimuli and a non-formal intuitive explanation like “In the verb-attached case the prepositional phrase says something about the verb”. Subjects were then asked to formulate the rule to distinguish the two attachments in their own words and were presented with four unambiguous practice items. Finally, they would read 80 to 100 sentences one by one and for each sentence decide between verb- or noun attachment. Ten seconds after a sentence appeared on screen a pop-up window encouraged subjects to answer faster. This time limit was chosen to force subjects to answer intuitively. However, many subjects would need more time on certain trials. After selecting their answer they could continue with the next item at their own pace. Half way through the questionnaire subjects were encouraged to take a longer break if needed. The stimulus list was split up into three parts to keep the duration of each survey to about 30 minutes. Each subject saw one of the possible lists in a pseudo-random order, so that sentences from the same pair were at least four items apart. Three subjects were excluded either based on poor performance on the practice items (less than three correct), because their average reaction time diverged extremely from the average (greater than 1.5 times the interquartile range) or because they had less than 60% correct answers to those sentences that were semantically completely unambiguous.

The survey results were analyzed using R version 3.6.3 and the lme4 package for linear mixed-effects models (Bates et al. [2015]). Pairs of sentences were selected if both received at least 74% of answers consistent with the intended attachment. With more than 74% of answers being consistent with the intended attachment we can exclude the alternative hypothesis of random behavior at an alpha level of 0.05 given a binomial distribution and 20 data samples per item. The selection was made so that every verb was repeated exactly two times and there were equal amounts of sentences in both verb and role condition.

On pre-test results for the final selection of sentences, we used a generalised linear mixed effects model (GLMM) with a logit link function fit by maximum likelihood to examine the relationship between accuracy (i.e. percentage of answers in line with our expectations), reaction time, plausibility ratings (on a scale of 1 to 5), context manipulation (verb condition or role condition) and attachment type (verb- or noun-attached). A mixed logit model appropriately accounts for binomial response variables (Jaeger [2008]), in our case hits or misses (correctly identifying an attachment according to intended sentence meaning or not). The model thus allowed us to test whether there were systematic differences in processing noun- or verb-attached sentences, as well as systematic differences between our different context manipulation conditions while controlling for between-subject variance. We specified accuracy (hit or miss) as the dependent variable and reaction time, plausibility rating, and context condition as fixed effects. Additionally, the model included random-effect terms for items (intercept only) and subject (intercept and slope). The model was fully saturated with all two-way interaction effects.

GLMM results indicate a significant effect of attachment type and plausibility, with factor level contrasts revealing that subjects were more often correct for noun-attached items (see Figure 3) and high plausibility ratings led to high accuracy. There was a significant Attachment type x Plausibility interaction. Factor level contrasts revealed that the effect of high plausibility leading to high accuracy was stronger for verb-attached sentences than noun-attached sentences (see Figure 4). The context manipulation effect was not significant and only the interaction Context Manipulation x Attachment was significant, indicating that only for noun-attached sentences were items more often correctly interpreted in the verb condition compared to the role condition (see Figure 3). Finally, the interaction of Reaction Time x Plausibility was significant. As illustrated in Figure 5, high plausibility ratings only lead to higher accuracy if reaction times were fast. In summary, whether sentences were constructed to fit the verb or the role condition did not lead to large differences in accuracies, although sentences in the verb condition were slightly biased towards a noun-attached interpretation. Most of the items used in the experiment received a plausibility rating of higher than 3 on average with only four items with an average rating below 3 and verb-attached sentences receiving on average slightly higher plausibility ratings.

**Figure 3:**
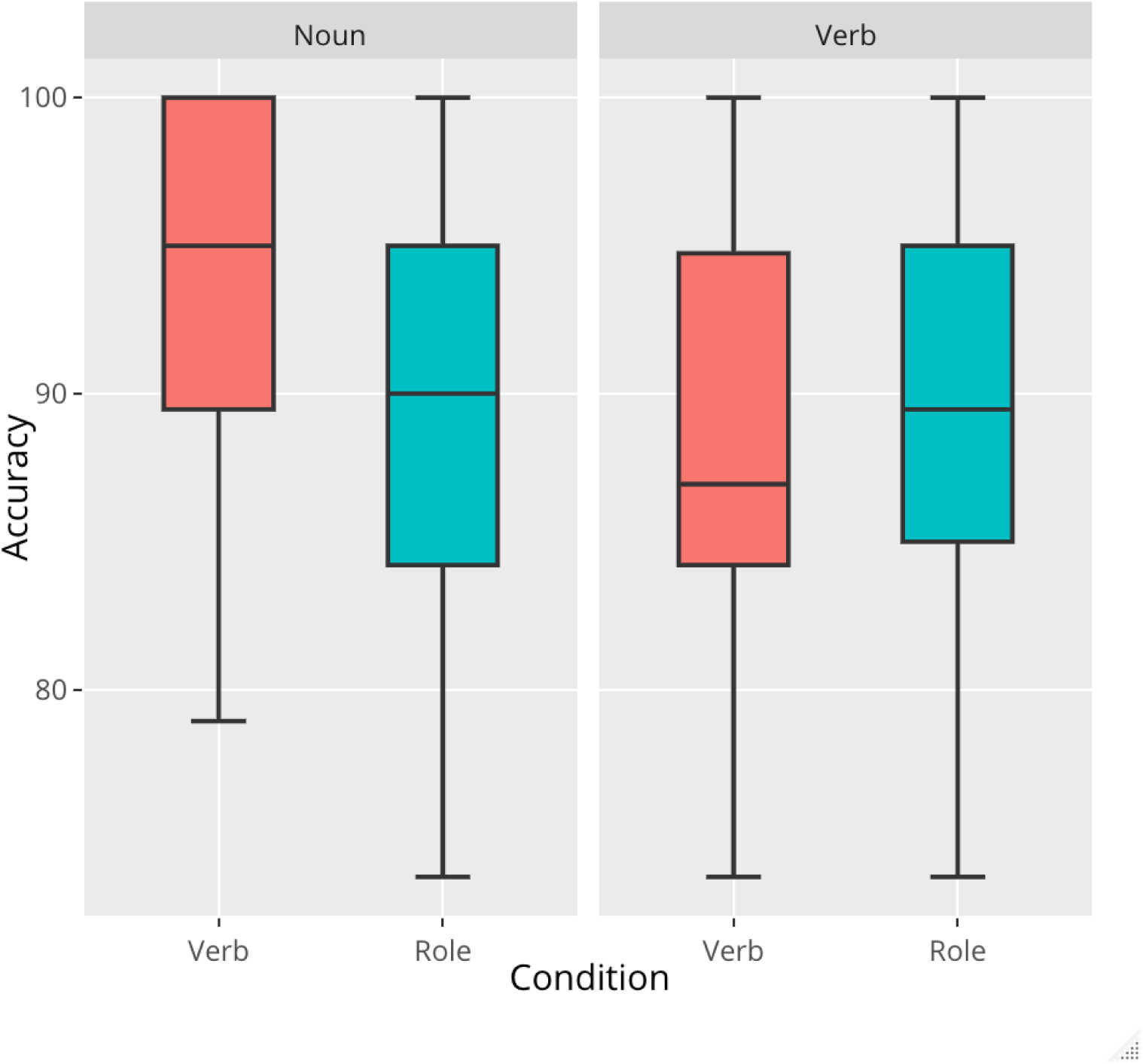
Pre-test proportion of correct responses averaged across all subjects. Accuracies are plotted separately for verb condition (red), role condition (blue) and for noun-attached sentences (leftmost graphs) and verb-attached sentences (rightmost graphs).

**Figure 4:**
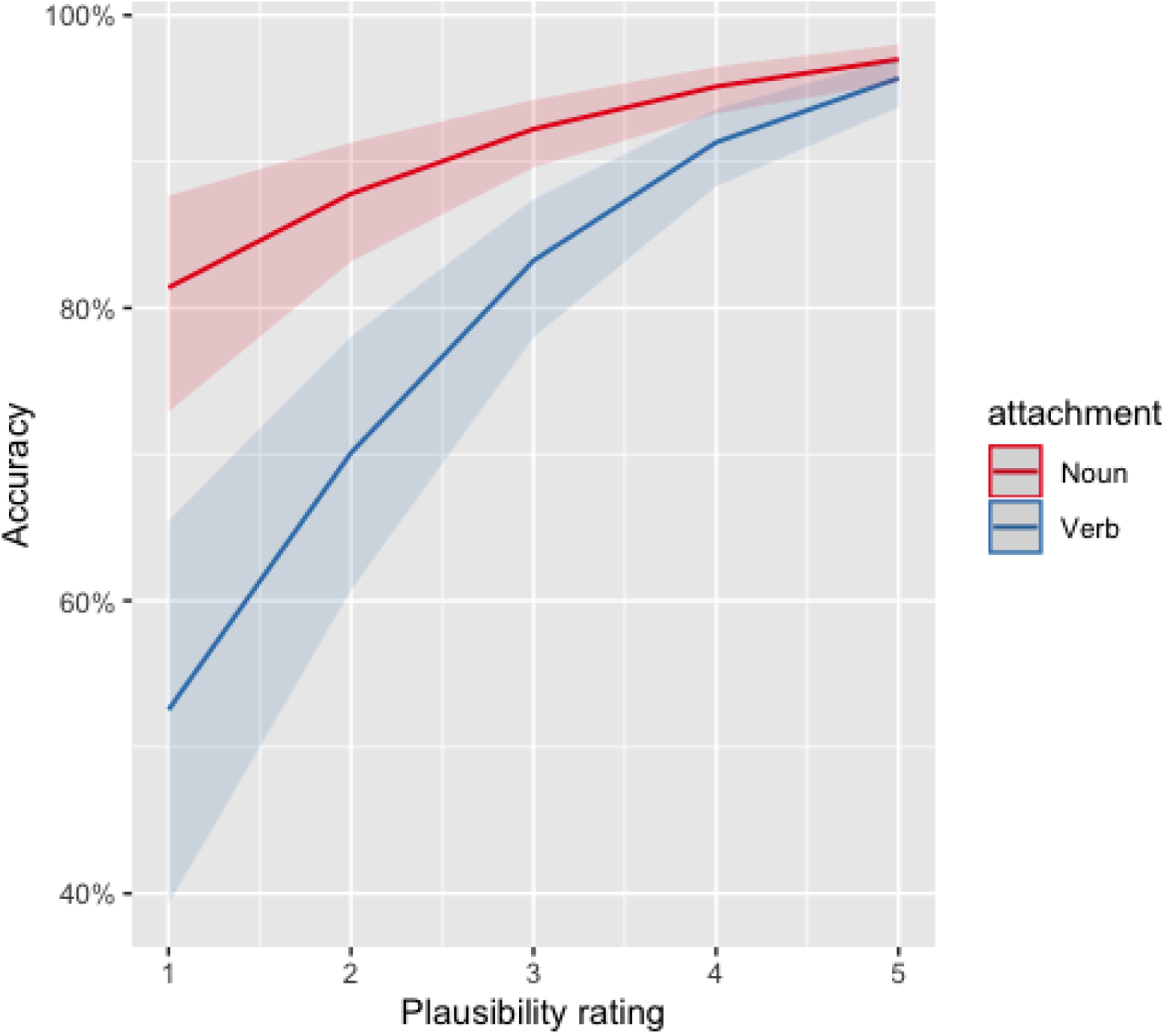
Interaction between plausibility ratings and attachment type. Mean Accuracy per plausibility rating is plotted for noun-attached (red) and verb-attached (blue) items.

**Figure 5:**
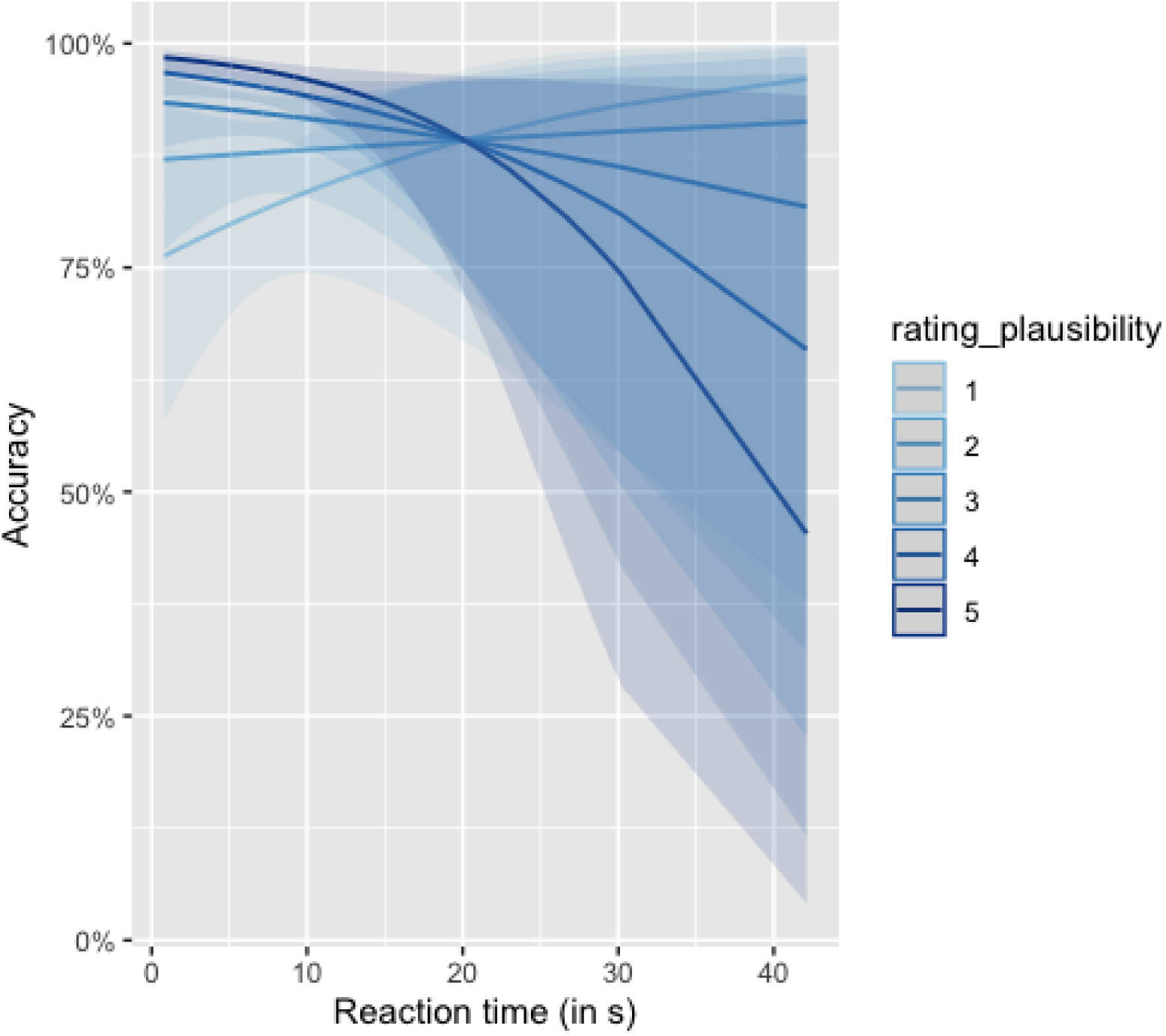
Interaction between reaction times and plausibility ratings. Mean Accuracy per reaction time is plotted for different plausibility ratings. The higher the plausibility the darker the color.

### 2.2. Experiment

10 Native German speakers (mean age = 22 years, 3 male) were seated in a magnetically shielded room and read sentences word-by-word while their neural activity was recorded using Magnetoencephalography (MEG). All subjects gave informed consent prior to filling in the survey and received financial reimbursement or credits. All stimuli were presented using the Presentation software (Version 16.0, Neurobehavioral Systems, Inc). Sentences were presented in pseudo-random order and word-by-word in four blocks with self-paced pauses in between blocks. In 25% of all trials a comprehension question would follow the sentence. Comprehension questions were either directed at identifying the agent or patient of the sentence (“Who has the bucket” or “Who is being carried”) or they would target the semantic dependency of the prepositional attachment (example question following (1): “Who has the questionable arguments”). The question was presented together with two answers, one on the left and one on the right side of the screen. Subjects indicated which answer they chose by pressing a button with their index finger corresponding to the position of the answer on the screen. The comprehension questions were meant to ensure that subjects were engaged and attentive during the task and that they fully parsed the presented sentences on both a semantic as well as structural level. Prior to the main experiment subjects received four practice trials to familiarise themselves with the pace of the presentation. Words were presented sequentially on a back-projection screen, placed in front of them (vertical refresh rate of 60 Hz) at the centre of the screen, in a white font, on a black background. Each word was separated by an empty screen for 200 ms and the final word of each sentence was followed by a 2000 ms blank screen. Duration of each word on screen was 392 ms on average and varied with word length with a minimum duration of 300 ms and maximum duration of 500 ms (formula: 300 ms + number of letters * 1000/60). The inter-sentence interval was jittered between 500 and 1000 ms. Within two weeks after the MEG experiment, subjects filled out a questionnaire rating each stimulus sentence as either noun- or verb attached and as plausible on a scale from 1 to 5. This questionnaire was the same as the one used for the pre-test but contained only those stimuli that had been used during the MEG experiment.

MEG data were collected with a 275 axial gradiometer system (CTF). The signals were analog low-pass-filtered at 300 Hz and digitized at a sampling frequency of 1,200 Hz. The position of the subject’s head was registered to the MEG-sensor array using three coils attached to the subject’s head (nasion, and left and right ear canals). Throughout the measurement, the head position was continuously monitored using custom software (Stolk et al. [2013]). During breaks the subject was instructed to reposition to the original position if needed. Subjects were able to maintain a head position within 5 mm of their original position. Three bipolar Ag/AgCl electrode pairs were used to measure the horizontal and vertical electrooculogram and the electrocardiogram. In addition to the brain signal, we acquired T1-weighted magnetic resonance (MR) images of each subject’s brain using 3 Tesla Siemens PrismaFit and Skyra scanners. All scans covered the entire brain and had a voxel size of 1×1×1mm^3. Finally, we recorded the subject’s head shape with the Polhemus for better co-registration of MEG and anatomical scans.

### 2.3. Preprocessing & Source reconstruction

Data were pre-processed using the Fieldtrip toolbox in MATLAB (Oostenveld et al. [2011]). For the decoding analysis the Donders machine learning toolbox (Van Gerven et al. [2013]) was used in combination with custom-made MATLAB scripts. The data were segmented into epochs around word onset with a 200 ms pre-stimulus period. To detect muscle artifacts, data was bandpass filtered between 110 Hz and 140 Hz and the trials with large variance were excluded upon inspection (less than 4% of all critical trials). Data was filtered between 0.1 Hz and 40 Hz. Independent component analysis (ICA) was used to remove artifacts stemming from the cardiac signal and eye blinks. For each subject, the time course of the independent components was correlated with the horizontal and vertical EOG signals as well as the ECG signal to identify and subsequently remove contaminating components.

We used linearly constrained minimum variance beamforming (LCMV) (Van Veen et al. [1997]) to reconstruct activity onto a parcellated cortically constrained source model. For this, we computed the covariance matrix between all MEG-sensor pairs as the average covariance matrix across the cleaned single trial covariance estimates. This covariance matrix was used in combination with the forward model, defined on a set of 7842 source locations per hemisphere on the subject-specific reconstruction of the cortical sheet to generate a set of spatial filters, one filter per dipole location. Individual cortical sheets were generated with the Freesurfer package (Dale et al. [1999],version 5.1) (surfer.nmr.mgh.harvard.edu). The forward model was computed using FieldTrip’s singleshell method (Nolte [2003]), where the required brain/skull boundary was obtained from the subject-specific T1-weighted anatomical images. We further reduced the dimensionality of the data, by grouping source points into 374 parcels, using a refined version of the Conte69 atlas. These parcels were used as searchlights in the subsequent analyses.

### 2.4. Multivariate decoding analysis

#### 2.4.1. Gaussian Naive Bayes

We trained a Gaussian Näıve Bayes classifier (GNB) (Mitchell [1997]) to identify cognitive states associated with underlying sentence structure from the pattern of brain activity evoked by reading the final word of a prepositional phrase. The GNB is a generative classifier that models the conditional probability *P* (*x_j_|Y_i_*) of signal amplitude x (at a given sensor/voxel j) given that the stimulus is of a class *Y_i_* (noun- or verb-attached prepositional phrase) using a univariate Gaussian and assuming class conditional independence. The mean and variance of this distribution is estimated on a subset of the trials (training set). The remaining data (test set) is then classified as the class *Y_i_* whose posterior probability *P* (*Y_i_|x*) is maximal among all classes. The corresponding classification rule is:

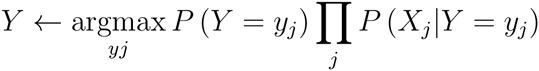

Classification results were evaluated using 20-fold cross-validation, so that accuracy was always based on test data that were disjoint from the training set. 20 folds were chosen for a good balance between amount of training data per fold and computational speed. Accuracy was estimated as the percentage of correctly classified trials across all folds. Classifiers were trained using a sliding time-window approach, where for each time-point, MEG data from all sensors and all time-points +-50ms were concatenated into a single vector (length = vertices x time-points). We also trained the same classifier on source-reconstructed data using a spatial searchlight approach in addition to the sliding time-window. The searchlight procedure followed the parcellation of the cortical sheet. For each parcel and time-point a classifier was trained on source data of all vertices within that parcel, while concatenating across all time-points within a sliding window of width 100 ms.

All parameters chosen for the classification analysis were manually optimised based on accuracy of an orthogonal classification task, namely to distinguish neural patterns evoked by either reading the main verb or the second noun (object noun) of the sentences. Decoding which of these different word classes was being presented robustly resulted in accuracies significantly higher than chance performance. Within our stimulus design, word class was confounded with ordinal word position in the sentences. Therefore, we conducted a control analysis on the same ordinal word positions within only filler items (where sentence structure varied and therefore nouns and verbs did not always occur at the same sentence position). This control analysis did not yield comparably high decoding accuracies. We compared the performance of the verb-noun classifier given different sliding time window widths (50 ms, 100 ms or 200 ms) and feature transformations (concatenating vs averaging over time dimension, feature selection, orthogonalisation and feature reduction through principal component analysis (PCA), gaussianisation).

PCA transforms the data into linearly uncorrelated components, ordered by the amount of variance explained by each component. Using these uncorrelated components as features can improve the decoding performance of classifiers such as GNB, which assume no feature covariance (Grootswagers et al. [2017]). Furthermore, PCA allowed our feature selection to be based on a data-driven approach by keeping only a subset of components that explain highest variance. We observed that both orthogonalising of features (sensor-time points) using PCA and feature reduction by restricting training to the first 60 components only, boosted classification accuracy. Further feature selection based on signal strength (selecting features based on largest difference in means between classes) did not improve accuracy beyond the the effects of feature reduction based on PCA. Gaussianisation of the sensor-level data prior to classification analysis or broadening the training time window did not yield large differences in performance. Based on these comparisons we then continued to train the classifier on the noun- vs. verb-attachments with the optimal parameters.

#### 2.4.2. Representational similarity analysis

Prepositional phrase attachment is interpreted based on the semantic information given the context preceding the phrase. We therefore predicted that there might be reactivation of this semantic information (i.e. those semantic features that most strongly influence the attachment) after the disambiguating sentence-final word. We tested this hypothesis through representational similarity analysis (RSA) (Kriegeskorte et al. [2008]), representing semantic content by means of a high-dimensional word-embedding vector (semantic vectors). For the word-embeddings we relied on pre-trained models published by facebookresearch^1^ which had been trained on German Wikipedia using fastText (Bojanowski et al. [2016]; Grave et al. [2018]).

First, we ensured that the semantic information captured by the word-embeddings is also encoded in the neural signal. We extracted all segments of neural data time-locked to each word presented and further restricted the selection to either content words only for this analysis or sentence-final words (as described in detail below). We then generated pairwise similarity measures between those words by computing the euclidean distance between their corresponding word-embedding vectors (semantic similarity model). Repeated presentations of the same word were treated as separate words (i.e. not averaged across). In the same way, we computed pairwise similarity measures for the corresponding segments in the neural signal, i.e. the pair-wise neural similarity during reading of the same words. Words that were not present in the vocabulary of the pre-trained embeddings were excluded from both semantic model and neural data, which left 387 trials in total. Neural similarity was computed based on a moving searchlight by concatenating all samples within a 100 ms time-window and across source locations within a given parcel, and this was repeated for all parcels and shifting time-windows (between word onset and 800 ms post onset) with an 80% overlap in time. Finally, semantic similarity and neural similarity were correlated (Spearman correlation) at each searchlight position. This resulted in a map indicating when and where neural activity reflected semantic information about the perceived words.

Crucially, we then generalised this RSA to the post-sentence phase, when subjects were reading the final, disambiguating word. For this, we re-computed the neural similarity, this time based on neural activity evoked by the final word. For each Verb-attached and each Noun-attached PP instead of the word-embedding of the final noun we assign the word-embedding vector of the preceding verb or noun respectively (i.e. of the most plausible attachment points). We then recomputed the euclidean distance between word-embedding vectors for all trial pairs, which now expresses for each sentence pair the semantics similarity with respect to the disambiguated attachment sites. Any significant correlations between the neural similarity and the attachment site semantic similarity indicate when and where neural patterns evoked by reading the final noun are also encoding (i.e. reactivate) information about the preceding verb or noun respectively.

### 2.5. Significance testing of decoding accuracy

When evaluating significance of group-level accuracy differences between two classifiers (GNB vs. logistic regression; part-of-speech classifier vs. word position control) we relied on non-parametric permutation testing (Maris and Oostenveld [2007]), randomly swapping observed accuracy between classifiers. For statistical evaluation of the GNB classifier against chance level we relied on information prevalence inference (Allefeld et al. [2016]) based on subsampling of single-subject permutations. Prevalence inference tests the significance of above-chance accuracy in the majority of subjects given the permutation distribution at an alpha level of 0.05. Permutation tests are preferred over traditional tests against theoretical chance level, given that the small amount of trials (typical for neuroimaging studies) will lead to larger cross-validation errors (Varoquaux [2017]). Therefore, we computed null-distributions on randomly re-labeled data for the GNB classification task. For the binary classification task we randomly selected half of the items per category (either attachment type of part of speech) and switched their labels in order to maintain an equal amount of items per class. For analyses conducted on the source-reconstructed data we used one fixed set of permutations of the observations for each searchlight to preserve spatial correlations.The procedure of generating a permutation and subsequent classification/prediction using permuted labels/semantic vectors was repeated 100 times per subject.

To evaluate statistical significance of the correlation values resulting from the RSA analysis, we used nonparametric permutation tests against a baseline of zero, including cluster-based correction for multiple comparisons across time and space.

## 3. Results

### 3.1. Behavioral

In the MEG experiment, all subjects had higher than chance level performance on answering the comprehension questions. On average they gave 77% correct answers on sentences from the verb condition, 72% correct answers for the role condition and 88% correct answers on filler sentences. While performance on the filler items was above chance for all subjects, some subjects performed at chance for questions from the verb and role conditions (see Figure 6). Since correct answers to target items depended on the interpretation of the prepositional phrase attachment, this suggests, that some subject’s attachment interpretations differed from the norm (as determined by the pre-test). Within a week after the MEG experiment, each subject had filled in an online post-test, explicitly rating all stimulus sentences as either noun or verb attached (following the methods from the pre-test). Average accuracy across subjects on this post-test did not differ between conditions (verb and role condition both 81% correct) and subjects interpreted the sentences mostly as intended. Except for two subjects, who performed close to chance, subjects had a minimum accuracy of 79% (see Figure 7).

**Figure 6:**
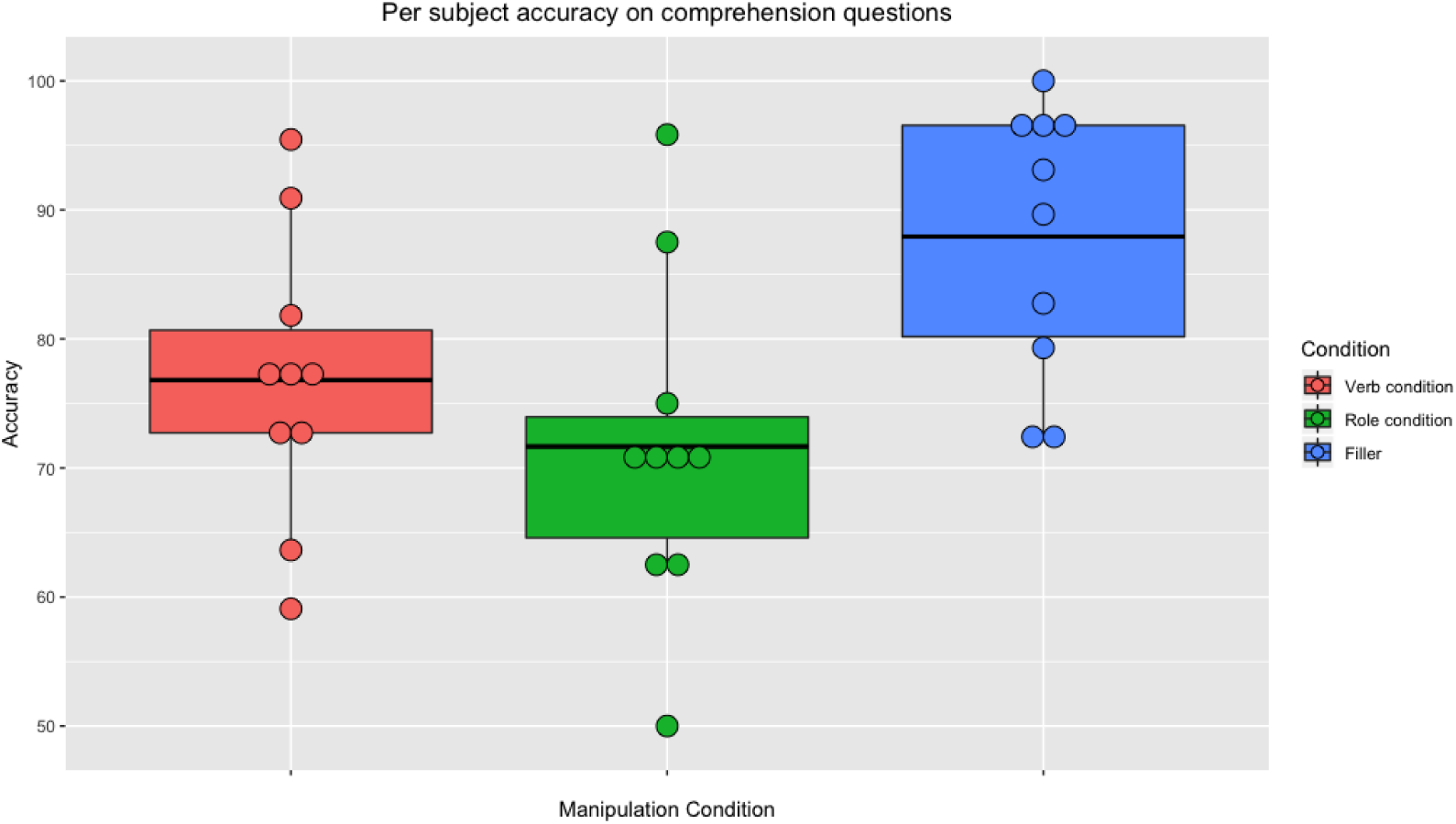
Accuracy of comprehension questions for each subject per manipulation condition. Accuracy across subjects depicted separately for each manipulation condition: Verb condition (left, red), role condition (middle, green) and filler items (right, blue). Individual subject accuracies are plotted as dots.

**Figure 7:**
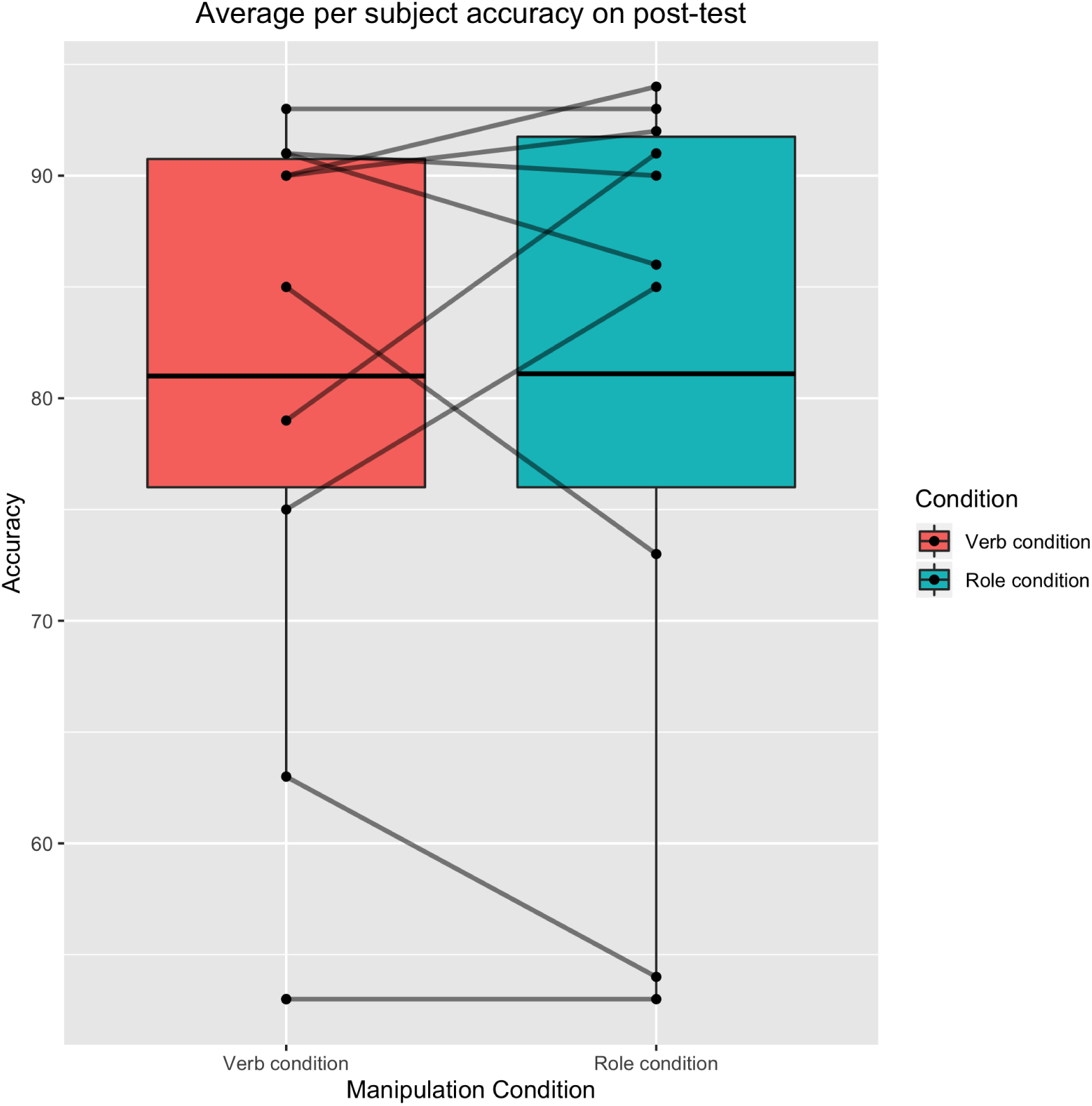
Accuracy of attachment rating for each subject per manipulation condition. Average accuracy is plotted separately for verb condition (left, red) and role condition (right, green). Individual subject accuracies (percentage of items correctly classified) are plotted as black dots.

### 3.2. Multivariate pattern analysis

#### 3.2.1. 2-way classification Noun-attached vs Verb-attached

Our main analysis of interest, the 2-way classification of different phrase structure (Noun attachment vs. Verb attachment) did not reach above chance-level accuracy at any time window up to 2 seconds after onset of the final word of a sentence. We observed this null-finding both, when items were labeled according to the general pre-tested attachments, but also when items were labeled according to subject-specific post-tests (see red and blue graphs respectively in Figure 8).

**Figure 8:**
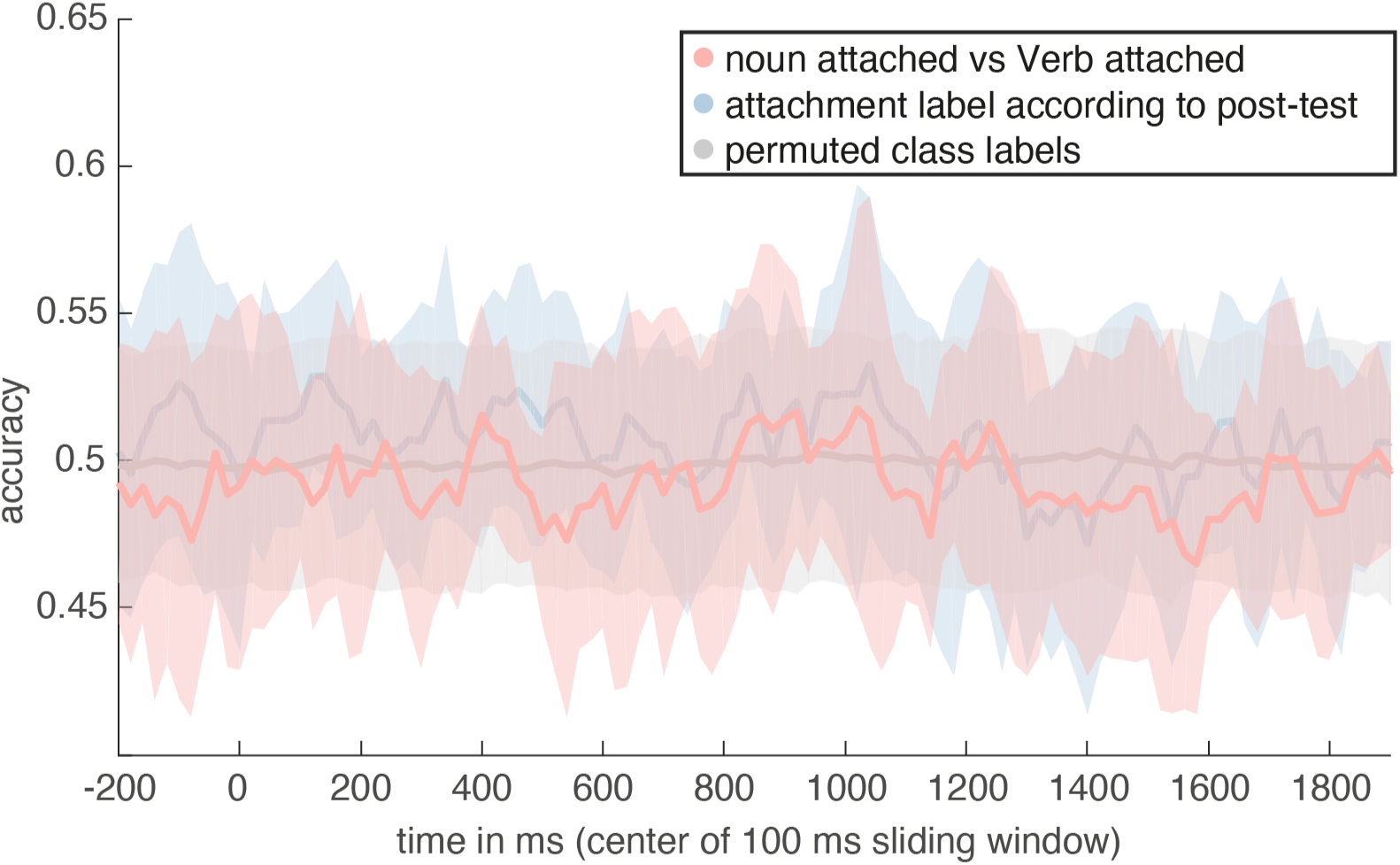
Attachment classification in sensor space. Accuracy of a Gaussian Naive Bayes classifier is plotted for two-way classification of attachment type (noun-attached vs verb-attached). Accuracy is shown for both, a classifier trained on items labeled according to coherent interpretations of sentences during pre-test (red) and a classifier trained on items labeled according to subject-specific post-test interpretations (blue). Observed accuracy was tested against a baseline performance estimate generated by repeatedly classifying data after permuting labels (grey).

#### 3.2.2. 2-way classification Noun vs Verb

The 2-way classification on whether the currently seen stimulus was a verb or a noun based on sensor-level MEG data reached a maximal average accuracy (across subjects) of 67% at 160 ms after word onset and was significantly more accurate as compared to the word position classifier (p=0, cluster-corrected permutation tests) up until 460 ms after word onset (see Figure 9). Note that classification accuracy is already significantly above chance before the onset of the noun/verb. This is due to the fact that nouns were always preceded by a determiner and verbs by a noun, effectively turning the baseline period into a determiner vs. noun classification sample. PCA transformation of the data led to higher classification accuracy as compared to training on the raw features. Additional feature selection based on class means did not lead to further increases in accuracy (see Figure 10). Training the classifier on moving windows of length 100 ms not only was more efficient in terms of computation time but also lead to higher classification accuracies as compared to training the classifier per time point (see Figure 11). Concatenating sensors of all time points mostly lead to slightly higher accuracies as compared to averaging over time points before training.

**Figure 9:**
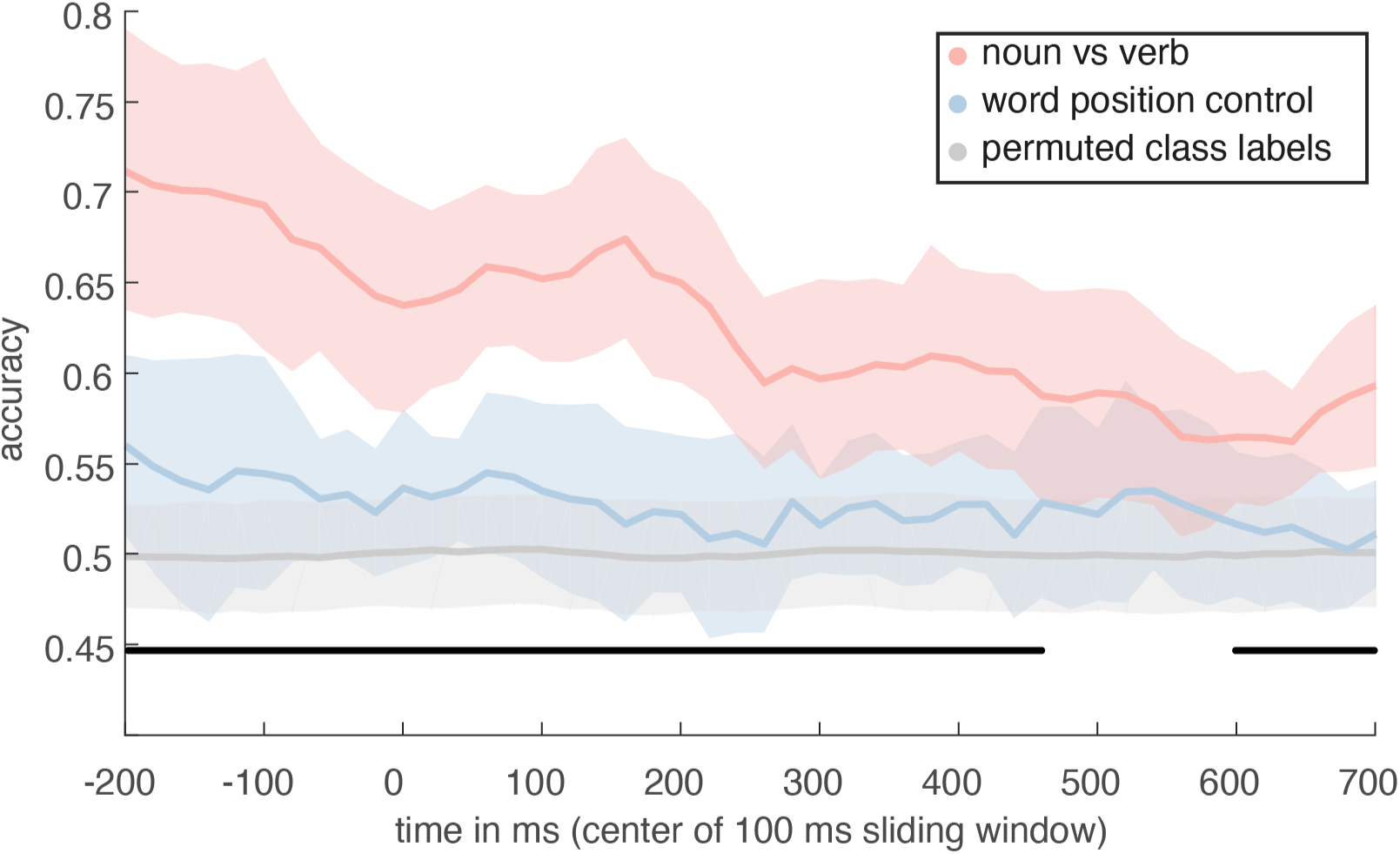
Part-of-speech 2-way classification in sensor space. Accuracy is plotted for part-of-speech classification (nouns vs verbs) using Gaussian Naive Bayes (red) and for classification of word position in filler sentences (blue, varying part-of-speech categories). Black lines indicate when part-of-speech classification is significantly higher as compared to classification on filler items. In addition, a chance performance distribution generated by repeatedly classifying data after permuting labels is depicted in grey.

**Figure 10:**
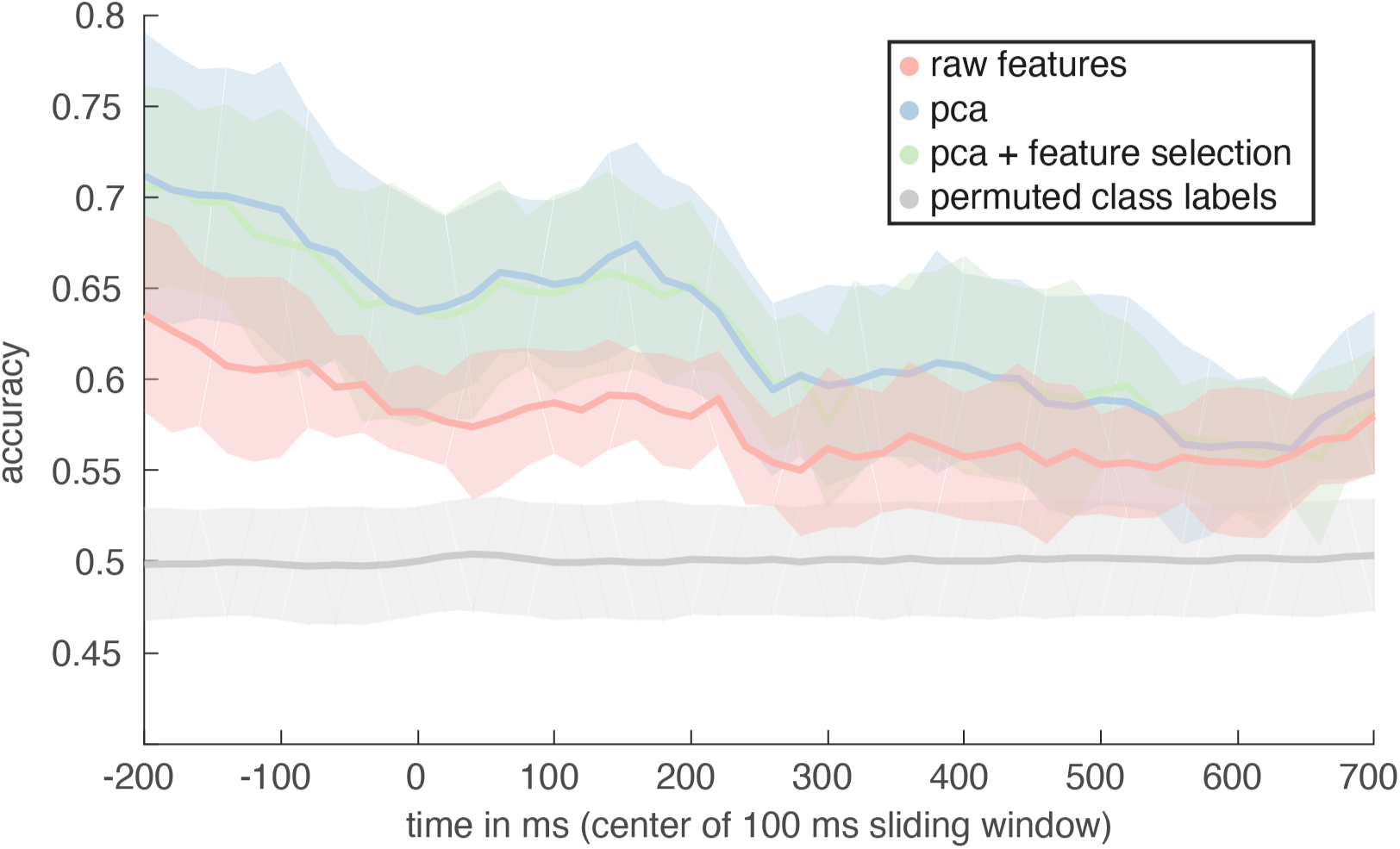
Feature transformation for 2-way classification. Accuracy is plotted for part-of-speech classification (nouns vs verbs) using Gaussian Naive Bayes and different feature reduction choices. Line plots represent the mean accuracy across all subjects and shaded areas represent its standard deviation. We first select evoked neural data from a 100ms (moving) time window and concatenate across all sensors and time points within that window, such that each sensor x time point equals one feature. We compare performance of a classifier trained on either the original features (red), on a dimensionality reduced sensor space after selecting only the first 60 components using principal component analysis (PCA, blue) or on a reduced feature space using PCA as well as further only selecting the 150 sensor x timepoints with the largest difference in class means (green). A baseline performance estimate was generated by repeatedly classifying data after permuting labels (grey). While feature space reduction through PCA improved classification accuracy, feature selection based on class means did not yield further improvements.

**Figure 11:**
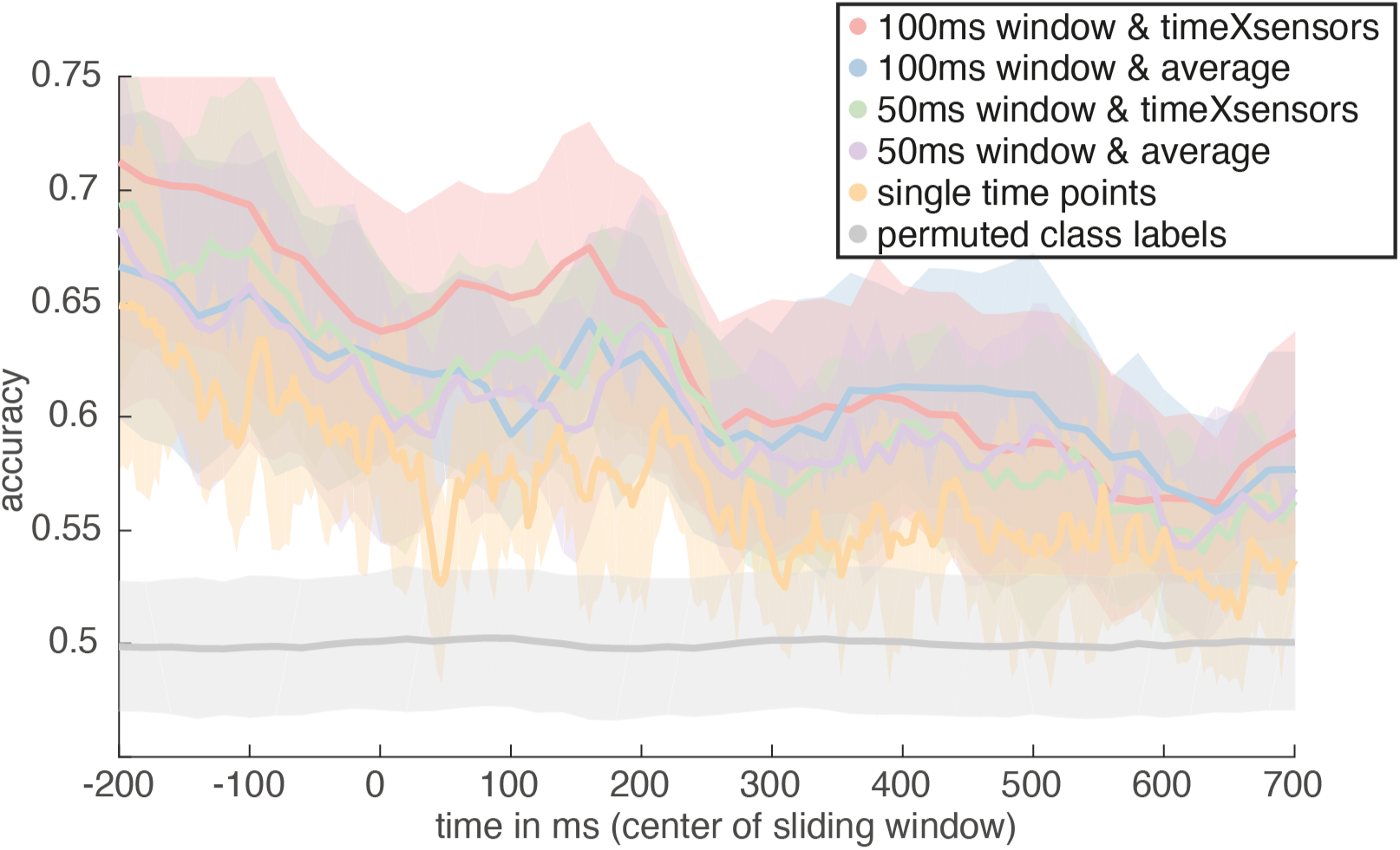
Time dimension for 2-way classification. Accuracy is plotted for part-of-speech classification (nouns vs verbs) using Gaussian Naive Bayes and different options for how to treat time. Line plots represent the mean accuracy across all subjects and shaded areas represent its standard deviation. A baseline performance estimate was generated by repeatedly classifying data after permuting labels (grey). Our moving window approach with window width of 100ms (red & blue) is most efficient in terms of computational time needed. On top of that, reducing the width of the window to 50 ms (green & purple) or even computing a separate model per time point (yellow) did not yield better classification performance. Further, for a window width of 100ms averaging over time points before training the classifier (blue) yielded lower accuracy as compared to concatenating across sensors and time points (red).

Besides Naive Bayes, we also tested different classification algorithms, i.e. support vector machines and logistic regression. None of these resulted in higher classification accuracies for the classification of nouns vs verbs (see Figure 12) as compared to Naive Bayes. Logistic regression performed better than Naive Bayes for the classification of determiner vs noun.

**Figure 12:**
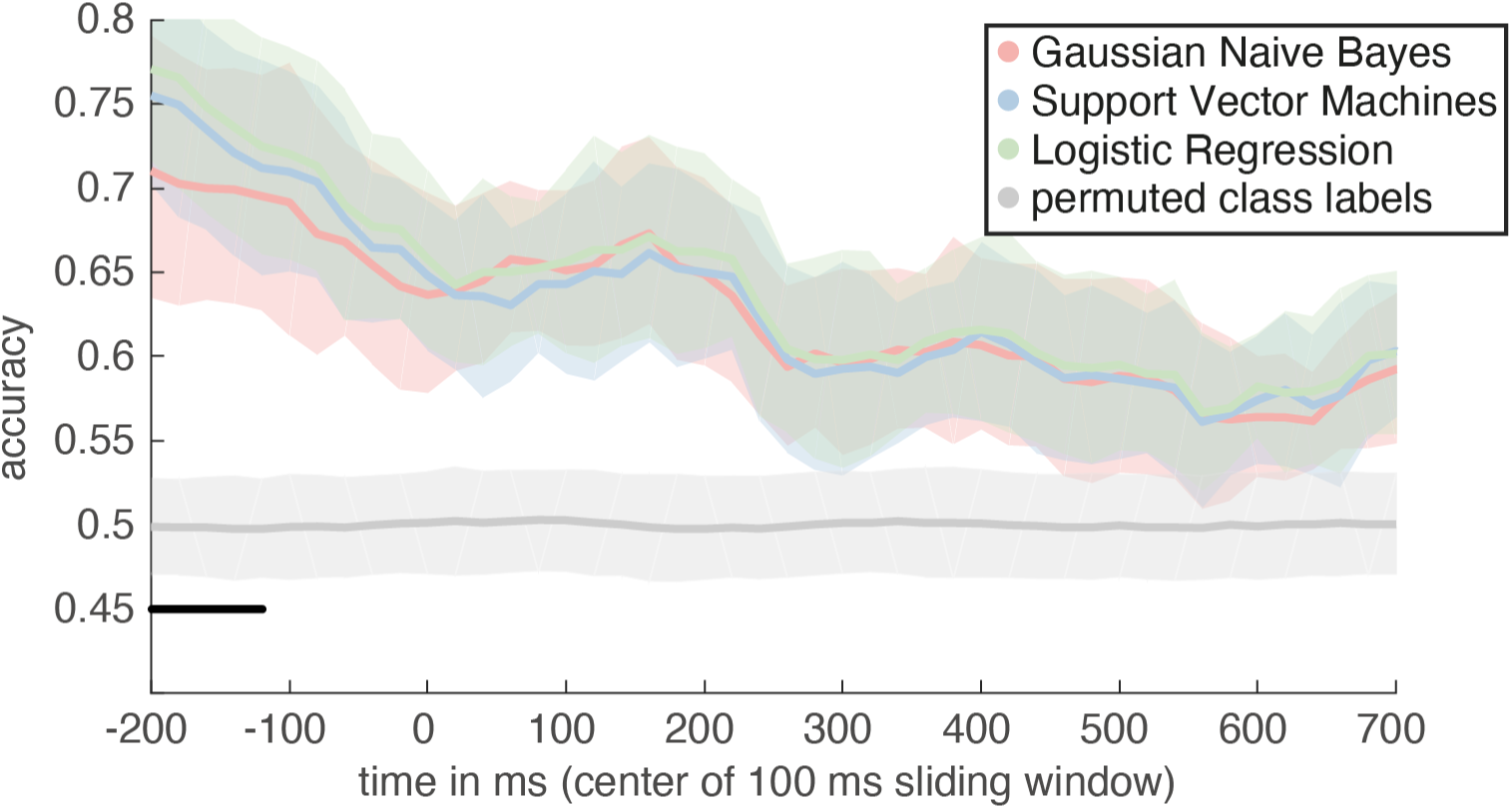
Comparison of different classification algorithms. Accuracy is plotted for part-of-speech classification (nouns vs verbs) using three different linear classifier: Gaussian Naive Bayes (red), support vector machines (blue) and logistic regression (green). Line plots represent the mean accuracy across all subjects and shaded areas represent its standard deviation. A baseline performance estimate was generated by repeatedly classifying data after permuting labels (grey). Significant differences in accuracy between different classifiers is indicated by a black bar.

Given that nouns and verbs have some systematic orthographical differences in German, we wanted to know whether classification success was mostly driven by low-level visual cortex. To investigate this, we source-reconstructed the MEG data and trained several classifiers on different regions across the cortex (searchlight approach). While classification accuracies were overall lower than those observed based on the sensor-level data, they were highest in occipital areas (see Figure 13). However, classification was also significantly above chance in more anterior cortical areas. With increasing time since word onset, classification accuracy increased as well in more anterior, bilateral occipito-temporal areas (see Figure 13 middle panel for Brodmann area 37). Between 340 ms and 540 ms, higher level areas like left inferior central and inferior frontal areas contain information about the noun-verb distinction (see Figure 13 lower panel for Brodmann area 43).

**Figure 13:**
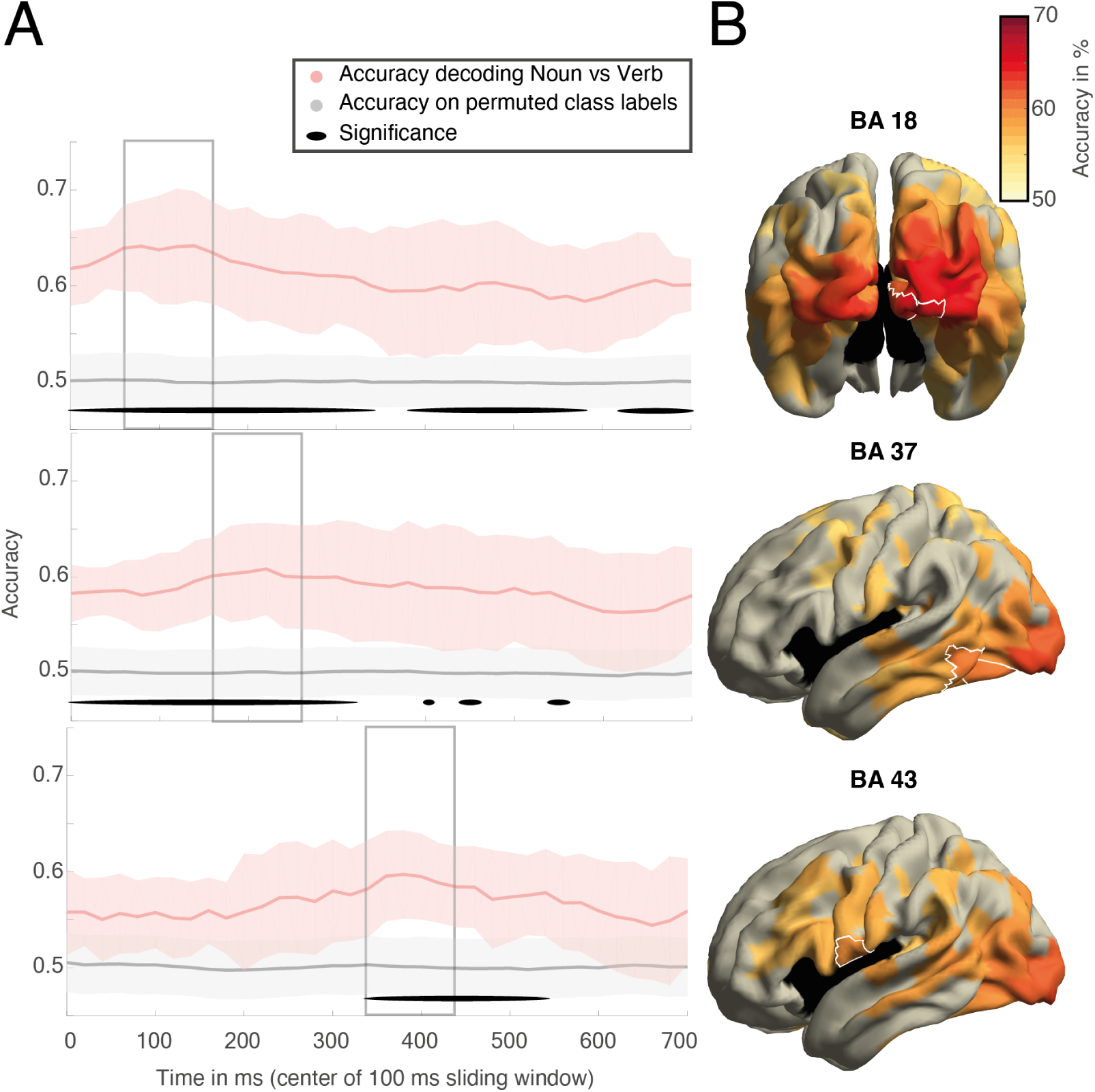
Part-of-speech 2-way classification in source space. Panel A: Accuracy is plotted over time for part-of-speech classification (nouns vs verbs) using Gaussian Naive Bayes (red). Observed accuracy was tested for significance (prevalence statistics, significant time points marked with black line) against a baseline performance estimate generated by repeatedly classifying data after permuting labels (grey). The upper, middle and lower panel display the mean accuracy over time for right occipital parcels (BA 18), left occipitotemporal parcels (BA 37) and left sub-central parcel (BA 43) respectively. Panel B: Cortical maps show the spatial patterns of classification accuracy, masked for significance. White contours outline the parcels for which time-courses are plotted in panel A respectively. Cortical maps contain averaged accuracies over the time-windows defined by the grey boxes.

#### 3.2.3. Generalization over time

Concerning the hypothesis that combinatorial processes involve a reanalysis of the to be combined parts, we tested whether after the onset of the final word of the sentence (the word which disambiguated the structural attachment of the prepositional phrase) the encoded information of the preceding noun or verb would be reactivated in the presence of either a noun- or verb-attachment respectively. We first investigated whether there was a reactivation of morphosyntactic information (part of speech) by generalising the 2-way classification trained on brain data measured during reading of noun and verbs preceding the prepositional phrase to the period following the final word of the sentence. Even though the final word was always a noun we hypothesised that only verb-attached prepositional phrases would in addition lead to verb-like activity patterns following the final word. However, contrary to our hypothesis the classifier trained on nouns and verbs in the context did not accurately classify the post-sentence period of verb-attached prepositional phrases as more verb-like (see Figure 14).

**Figure 14:**
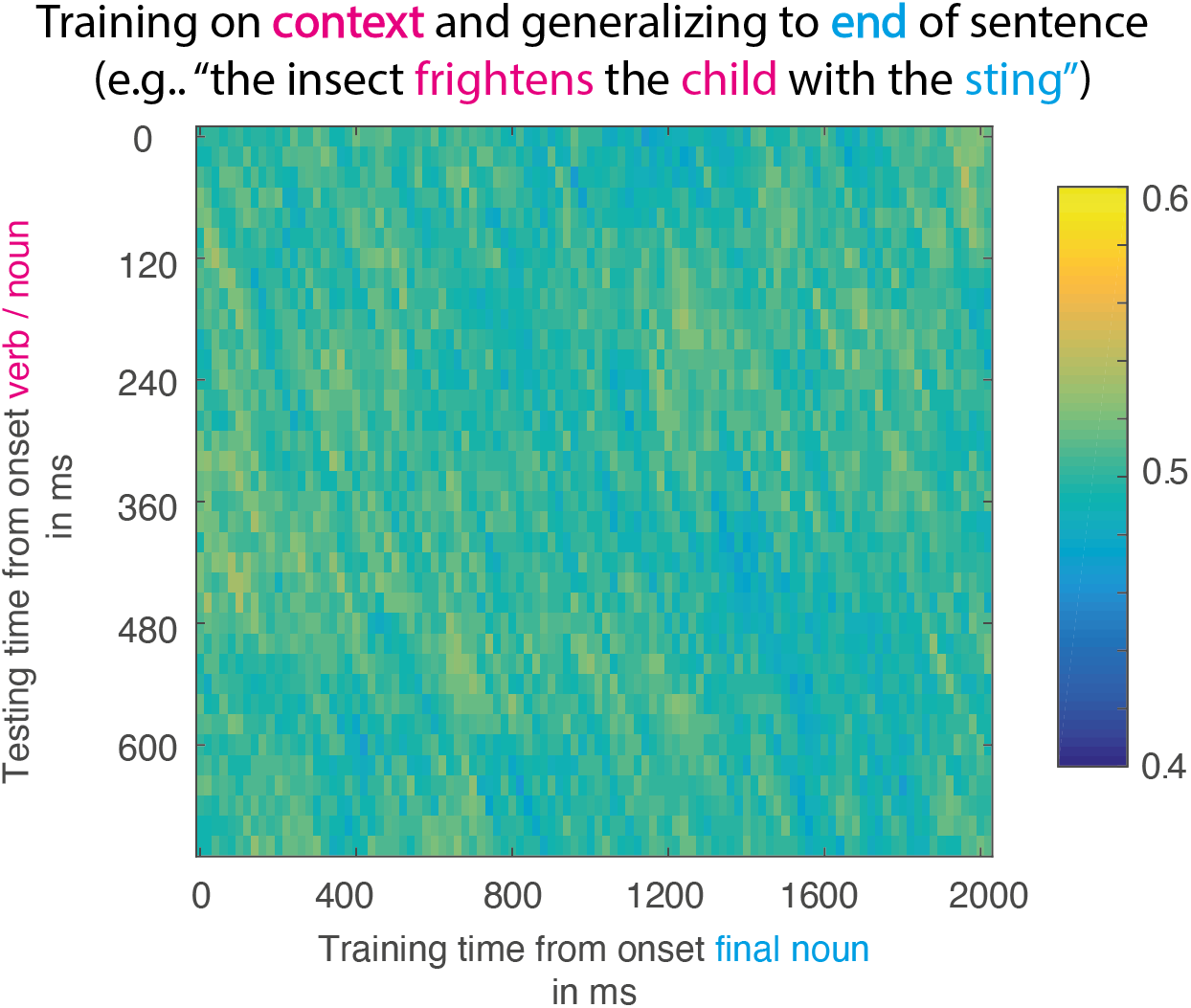
Generalised classification accuracy for part-of-speech. We trained a classifier to distinguish between nouns and verbs based on the neural data evoked by reading one or the other. While training this classifier on a moving time window starting at onset of the noun/verb, we then tested whether the learned weights would generalise to data recorded while reading the end of the corresponding sentence. To illustrate this on a specific stimulus example, on a sentence like 7 “The insect frightens the child with the poisonous sting”, we would train the classifier on distinguishing activity evoked by “frightens” from activity evoked by “child” but we would test the classifier on activity evoked by “sting”. Given that this sentence contains a verb-attached preposition, the correct label for the classifier to identify would be “verb”, regardless of the final word always being a noun. Color codes for classification accuracy at any given training-by-testing time tile. Generalised classification accuracy is not significantly above chance-level at any time point.

#### 3.2.4. RSA

For our stimuli, the interpretation of a prepositional phrase attachment was purely driven by semantic content. Therefore, we might also expect any reactivation to occur in the form of semantic information. We therefore tested whether at the time of disambiguation, any of the semantic information of preceding context would be reactivated. Specifically, we expected the semantics of the verb to be more strongly activated at the end of a verb-attached prepositional phrase and the semantics of the noun to be more strongly activated at the end of a noun-attached prepositional phrase.

Our RSA revealed significant correlations between a model of the trial-by-trial similarity derived from word embeddings and the pairwise similarity derived from neural data evoked by the corresponding words (see Figure 15). Activity patterns that correlated with semantic similarity first emerged in a window from 380 ms to 480 ms in superior parietal cortex. Between 440 and 600 ms after word onset, semantic information was represented more extensively across parietal, temporal and occipital regions. Areas in which activity patterns significantly correlated with semantic similarity included posterior parietal cortex, somatosensory cortex, angular gyrus, fusiform gyrus, auditory cortex and posterior parts of the superior temporal gyrus. Late after onset, from 560ms to 720ms only areas in the ventral occipital lobe remained significantly correlated. When we generalised the RSA to the final word of the sentence, however, there was no significant correlation with semantic similarity in any brain area and hence no evidence for semantic reactivation.

**Figure 15:**
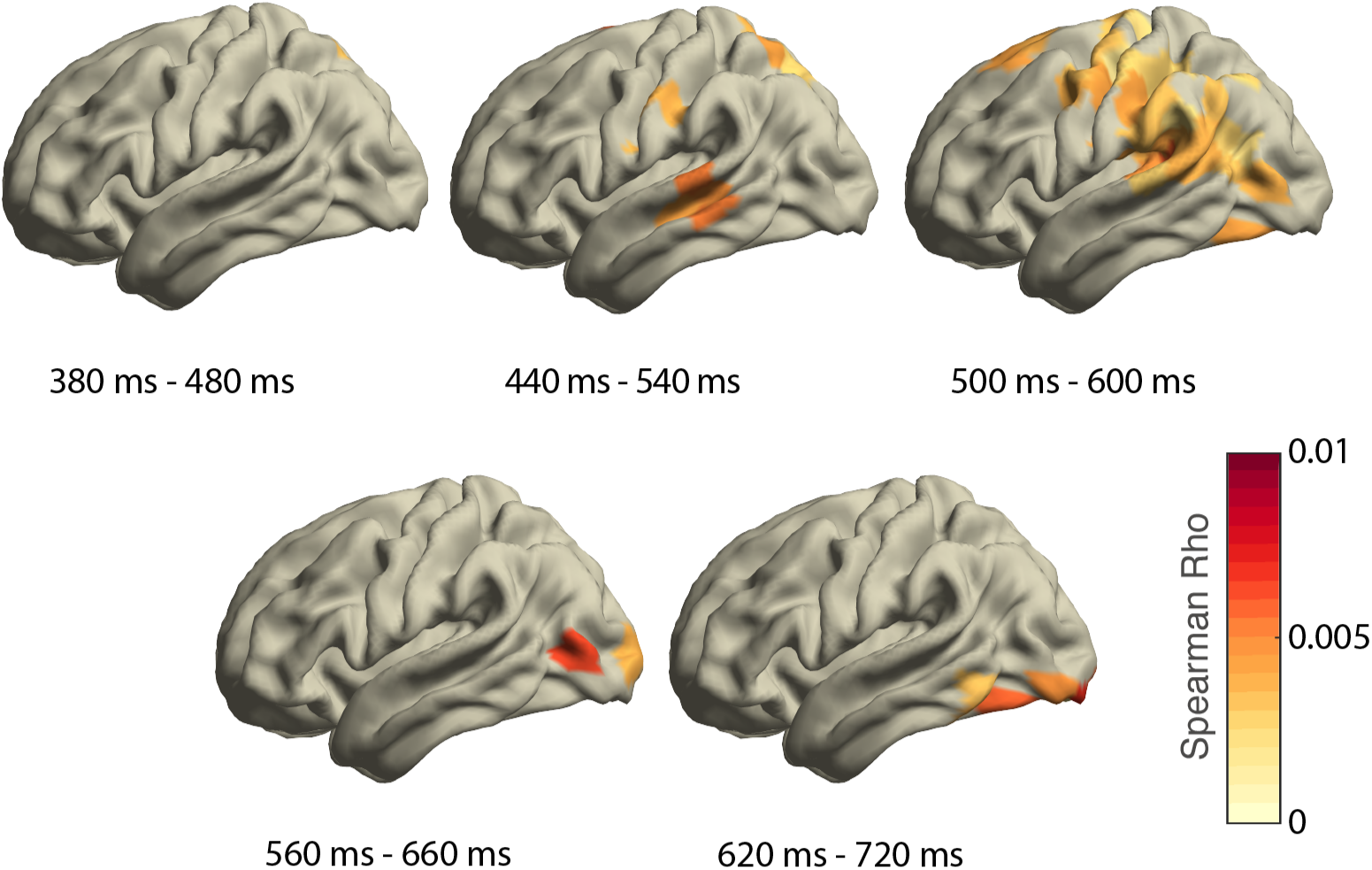
Searchlight RSA analysis on semantic information as measured by word embeddings. Cortical maps show the spatial patterns of correlations with the semantic similarity model (masked for significance) averaged across several time windows. Colour codes strength of correlation.

## 4. Discussion

In this study we applied MVPA to probe the neural signal for hierarchical structure building during online reading of structurally ambiguous sentences. Subjects read sentences containing verb-attached and noun-attached prepositional phrases ambiguous with respect to their attachment. We successfully applied a Naive Bayes classifier to classify part-of-speech information of the current stimulus from the multidimensional evoked neural activity. We also successfully extracted neural patterns encoding semantic information of content words as subjects were reading them, through modelling the pairwise semantic similarity structure of all word pairs (RSA) with corpus-extracted word-embeddings. However, none of these measures revealed encoding of different underlying hierarchical phrase structure for verb- vs noun-attached sentences at the end of the sentence, when attachment information was disambiguated through combined semantic information. That is, we did not find traces of stronger reactivation of either verb or noun in verb- or noun-attached sentences respectively; not in terms of their part-of-speech identity nor in terms of their semantic content. Nor were we able to directly train a classifier to distinguish between verb- and noun-attached PPs across varying lexical material. In the following, we will discus several potential explanations for the absence of an effect.

### 4.1. Signal-to-noise ratio

Could it be that our analyses were simply not sensitive enough to reveal effects of high-level processes such as phrase structure building? Previous literature relying on MVPA to capture higher-level language processing does not necessarily suggest high-level effects to be smaller as compared to more perception related effects. For example, Tyler et al. used an RSA approach to investigate the temporally unfolding syntactic computations during listening of temporarily ambiguous sentences (Tyler et al. [2013]). While their more perceptual word identity model correlated robustly with neural activity (rho *>* 0.015), when probing more abstract syntactic processing they found both small and large effects. Specifically, their model quantifying verb subcategorization information was only marginally significant and correlations were much weaker (rho *≈* 0.005) and only occurred on the word following the verb (n+1). Their model distinguishing ambiguous from unambiguous sentences, however, correlated even more strongly (rho *>* 0.020) with neural activity at late time points. Unfortunately, it is not straightforward to compare these effect sizes to our study. Our approach is novel in that we tried to directly probe neural representations of hierarchical phrase structure rather than its consequence on ongoing processing demands (e.g. memory requirements Nelson et al. [2017] or processing effort due to ambiguity Tyler et al. [2013]). Therefore, it is not immediately clear from those prior studies whether an MVPA approach is powerful enough to reveal representations of phrase structure directly.

Through additional analyses, targeting orthogonal syntactic information such as part-of-speech we tried to somewhat assess the sensitivity of our approach. Our Naive Bayes classifier reached a maximum average accuracy of 67% when trained to distinguish nouns from verbs. Above chance level performance was observed robustly across all subjects. Part-of-speech although not directly indicative of hierarchical structure, is a higher-level syntactic feature and hence our classifier captured information beyond perceptual signals. It is important to note, that within our design, the part-of-speech contrast is partly confounded by physical attributes of the stimulus. Specifically, nouns and verbs differ in their form as well as their syntactic function (e.g. the majority of verbs ended in the same inflexional syllable -t signalling third person singular). We must assume that any decoding success is partly due to stimulus form. Still, our observations that part-of-speech information can be decoded from anterior brain regions in addition to occipital cortex suggests that information was not solely based on the wordform differences. Hence, while the part-of-speech classifier provides some indication to the utility of the data with respect to higher-level features, it does not necessarily ensure the success of decoding more higher-level phenomena such as hierarchical structure.

Furthermore, we also set out to find semantic and syntactic reactivation of structurally relevant context as a direct consequence of phrase structure building. Brain data and semantic models correlated with a maximum correlation coefficient smaller than 0.01. This coefficient describes the correlation with data evoked by stimuli on screen and correlations can be expected to be substantially smaller when looking at the reactivation period. It is plausible to assume that reactivated neural patterns are harder to detect, as they are not directly evoked by a stimulus. In the present analyses, we focused on the time window following the onset of the final word. Content of the final word, however, was orthogonal to the supposedly reactivated information. For example, the last word of the sentence was always a noun and the same nouns (same semantic information) were presented in both verb- and noun-attached version. Nonetheless, in half of the trials (namely the verb-attached phrases), we would expect reactivation to reflect semantic and syntactic information of the preceding verb. The question is, whether MVPA is sensitive to internally generated, behaviourally relevant information, even with interfering material driving the neural response. While decoding of semantic category membership has been shown in the absence of a stimulus on screen (Simanova et al. [2015]), this was only shown for single words. To our knowledge there are no language studies explicitly probing reactivation in sentence context through MVPA. Within vision research, however, it has been shown that during a visual working memory tasks, information about stimulus orientation could be decoded from EEG during the retention period only through perturbation using an impulse stimulus (so called ‘ping’) but would otherwise be undetected (Wolff et al. [2017]). The authors argue that relevant information is not encoded explicitly in a persistent activity state but through an item-specific neural response profile that needs to be probed in order to affect ongoing neural activity. This might also explain why previous effects of prepositional phrase attachment ambiguity were found not directly following the disambiguating word but on subsequent words (Taraban and McClelland [1988]; Boudewyn et al. [2014]). Since we did not have a sentence continuation after the disambiguating noun, we may have been less sensitive to alterations in response profile caused by attachment structure.

Finally, it is possible, that our sensitivity was reduced by temporal variability in processing of the ambiguous sentences. It can be observed in the literature, that decoding accuracies are usually largest soon after stimulus onset and then decrease with increasing time (Cichy et al. [2014]; van Es et al. [2020]). We observe a similar pattern for our part-of-speech classification performance, which peaks very early after word onset (160 ms) but then decreases sharply until 250 ms after onset and continues to decrease thereafter. Thus, most information seems to be already encoded in the onset-potential or at least the neural signal might become more salient due to onset-related synchronisation of postsynaptic potentials. Effects of hierarchical structure building however may be less strictly time-locked events. Specifically, the varying difficulty in resolving structural ambiguities in our stimuli might have caused the signal to be jittered in time such that any reactivation might be less consistently synchronised across trials and subjects. Generally, each stimulus evokes a cascade of brain processes (both bottom-up and top-down) which all can vary slightly in their duration depending on context and individual and may therefore lead to more substantial variation in later, high-level brain processing as compared to initial bottom-up processing. Such temporal variability might have led to lower sensitivity for finding our effect as well. Future analyses should take temporal variability explicitly into account to not encounter the same issue. To achieve this, probabilistic frameworks for data-driven estimation of brain states could be used to align processing and overcome temporal variability. For example, Vidaurre et al. have developed an analysis that not only defines multiple representational states that dynamically encode the stimulus but also specifies which of these states is active when in time (Vidaurre et al. [2019]).

### 4.2. Shallow processing

Assuming that our signal to noise ratio in principle allows to capture neural representations of hierarchical structure, we will now turn to some more cognitive explanations for our failure to decode such structural representations. It is possible that readers do not compute phrase structure by default and at all times. Specifically, our experiment may have discouraged any detailed syntactic processing and subjects may have been engaged in “shallow” processing instead, similar to what has been reported before for garden-path sentences under the term “good-enough processing” (Ferreira and Patson [2007]; Ferreira and Lowder [2016]; Traxler [2014]). The idea of good-enough processing is that readers often arrive at a semantic proposition when interpreting a sentence without conducting a full syntactic (re)analysis. The recently established link between shallow processing and information structure (Ferreira and Lowder [2016]) further increases the plausibility of prepositional phrases falling victim to this strategy as well. Specifically, Ferreira & Lowder suggest that processing effort is usually directed towards parts of a sentence that constitute new rather than given information. The motivation for such a strategy is twofold. Firstly, it would maximise the success of integration of newly received information. And secondly, since given information links to prior discourse it is also more likely to be redundant and therefore more likely to survive “shallow” processing. It might not be obvious why our experiment should be affected by such shallow processing, given that we presented subjects with unrelated sentences without any larger discourse context to drive information structure. PPs are, however, making up the subordinate clause of the sentence, which is standardly viewed as communicating previously known information (Hornby [1974]) rather than new. Hence, it is possible that structurally inherent information structure in sentences with PPs causes readers to allocate less processing resources onto the structural disambiguation of the attachment. This would also be in line with processing accounts where hierarchical operations are not assumed as the default (Frank et al. [2012]). It is assumed that such processing strategies can be overwritten by strong task demands. For example, previous research has shown that syntactic task demands can reveal a P600 when there was none evoked by a purely semantic task (Mongelli [2020]). Indeed, many previous studies probing syntactic processing make use of syntactic tasks such as grammaticality judgments (Tyler et al. [2013]). In our study, however, subjects had to respond in only 25% of the trials and even on those trials, comprehension questions were not always probing knowledge about the PP region. The absence of a task and the fact that thematic role assignment could only be based on semantic cues in the first place may have discouraged a deep analysis of phrase structure.

The good-enough processing hypothesis further implies that hierarchical structure need not be computed at all in order to assign thematic roles. Instead, the semantic implications of the assigned thematic roles would be the sole outcome of successful sentence processing. Semantics of thematic roles are more complex and numerous than their possible corresponding phrase structures. Through adopting a strictly binary distinction of verb- and noun attachments we have intentionally ignored this semantic variation to target only the structural differences. However, as mentioned before, phrase structure and thematic roles are somewhat related and hence can easily become confounded. In fact, the relationship between thematic roles and syntactic structure is somewhat asymmetric to begin with. While any given thematic role is always bound to a certain syntactic structure^2^, this is not a bidirectional relationship. For example an instrument role will always be expressed in a verb-attached PP, but not every verb-attached phrase structure is necessarily carrying information about instruments (see sentences 8 & 9 for alternative role example).

8. The girl cuts the apple with a knife. (instrument role)

9. The girl cuts the apple with vigour. (manner role)

Taraban et al. have shown that previously reported reading time effects of PPs can be explained largely by expectations about thematic role. Specifically, they showed that unexpected structural attachment (verb- or noun attachment) do not delay reading times beyond the effect of thematic role expectations (Taraban and McClelland [1988]). The P600 effects reported by Boudewyn et al. could have also been driven by the semantics of the associated thematic roles rather than structure per se. In their stimulus set all verb-attached stimuli contained PPs expressing an instrument and all noun-attached PPs expressed an attribute. Moreover, most of their sentences contained action verbs (which bias towards expectations for instrument roles to begin with). Their P600 could therefore just as well be a marker for surprisal due to the unexpected thematic role in noun-attached sentences. In our study, we had more varying verb types (almost a third of all verbs were perception verbs) and more varying thematic roles (see table 1). However, the definition of thematic roles can be murky and the less common ones are usually poorly defined. With the exception of agent and patient role, the psychological reality of certain thematic roles (even as prominent as the instrument role) can be debated (Rissman and Majid [2019]). It is therefore difficult to systematically manipulate this dimension. Nonetheless, through using more varied thematic roles and verbs we have created a more naturalistic stimulus set as compared to previous studies, potentially weakening effects of thematic role expectations, that likely have been driving previous findings of divergent neural activity between noun- and verb-attached PPs.

In conclusion, with this study we could not identify a neural representation of hierarchical structure using MVPA. We did show, however, that our MVPA approach was in principle sensitive to both syntactic and semantic information encoded in the neural signal. Further, we did not find any differences between processing verb- or noun-attached prepositional phrases unlike previous studies have suggested. We speculate that this was partly due to our well controlled and semantically varied sentence material. In the future, a more fine-grained characterisation of the semantic dimensions driving attachment decisions and the systematical manipulation of thematic roles may help to establish any differences in processing PPs at a purely structural level.

## 6. Appendix

### TIGERSearch queries

We defined the number of ambiguous prepositional phrases (PPs) as those phrases that dominate a preposition and directly follow a noun:

(1) [pos=“NN”].#pp:[cat=“PP”]& #pp *>* #prep:[“APPR” | pos=“APPRART”]

We extracted frequency counts for all postnominal modifiers (noun-attached) within the ambiguous PPs, excluding those cases where the PP is topicalized (sentence-initial and therefore not ambiguous):

(2) #noun:[pos=“NN”].#pp:[cat=“PP”]& #phrase *>* #noun & #pp *>* #prep:[pos=“APPR” | pos=“APPRART”]& #n *>*MNR #pp & #phrase *>* **?** #x & [cat=“VROOT”] !*>***?** #x

Similarly, we extracted frequency counts for all verb modifiers (verb-attached) within the ambiguous PPs:

(3) #noun:[pos=“NN”].#pp:[cat=“PP”]& #pp *>* #prep:[pos=“APPR” | pos=“APPRART”] & #n *>* MO #pp

### Stimulus Material - Verb condition

**Table 2:**
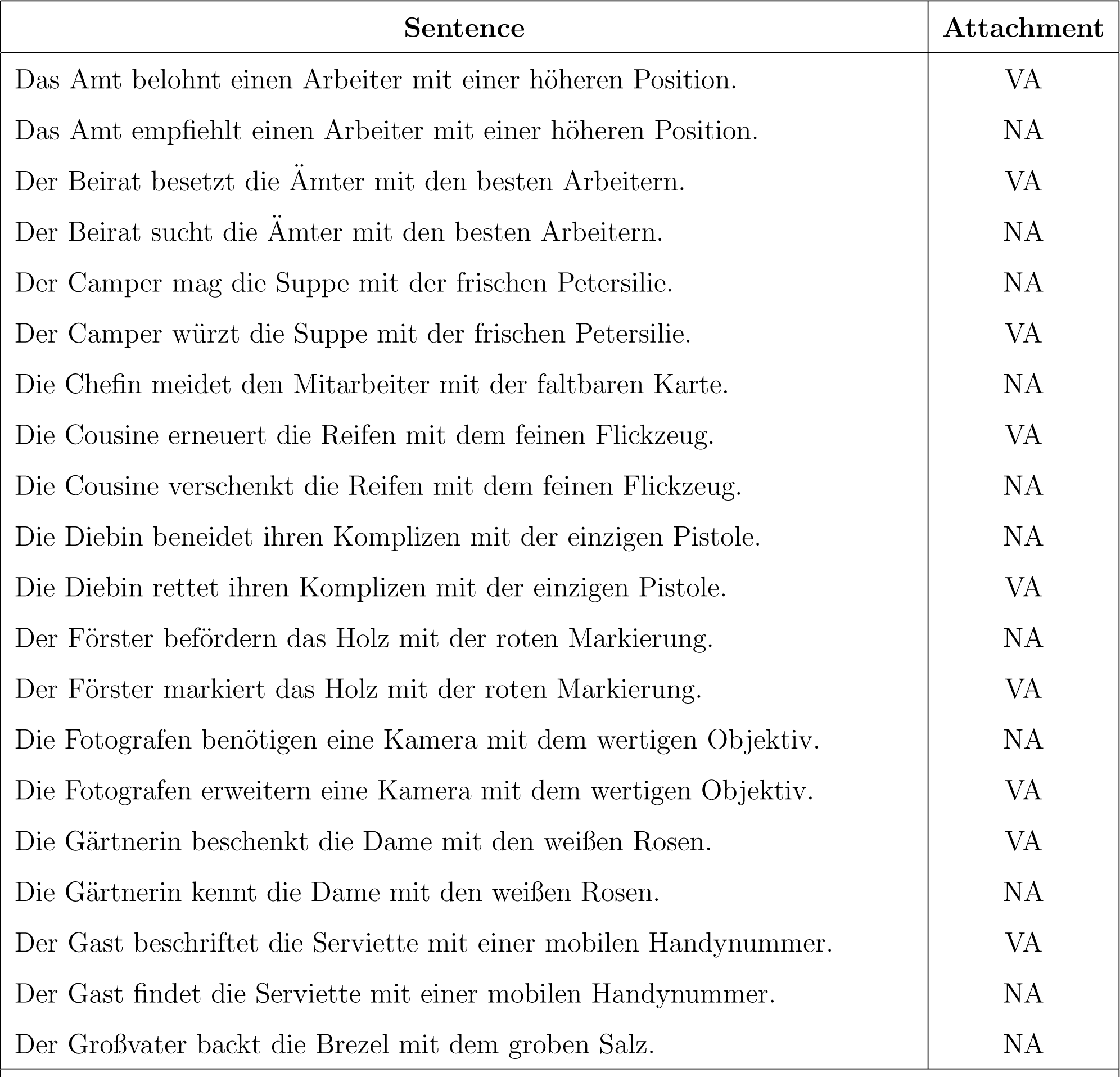
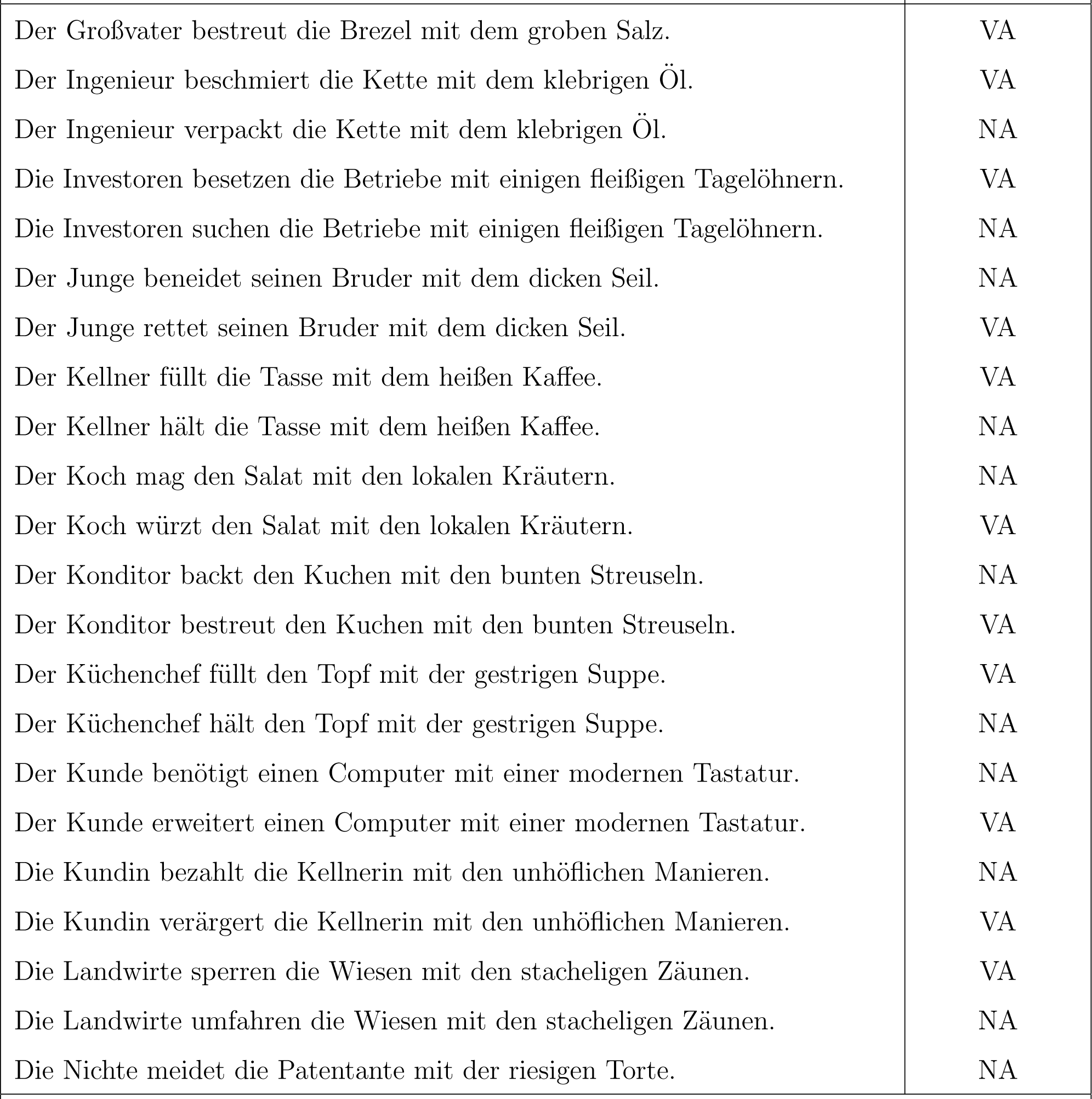
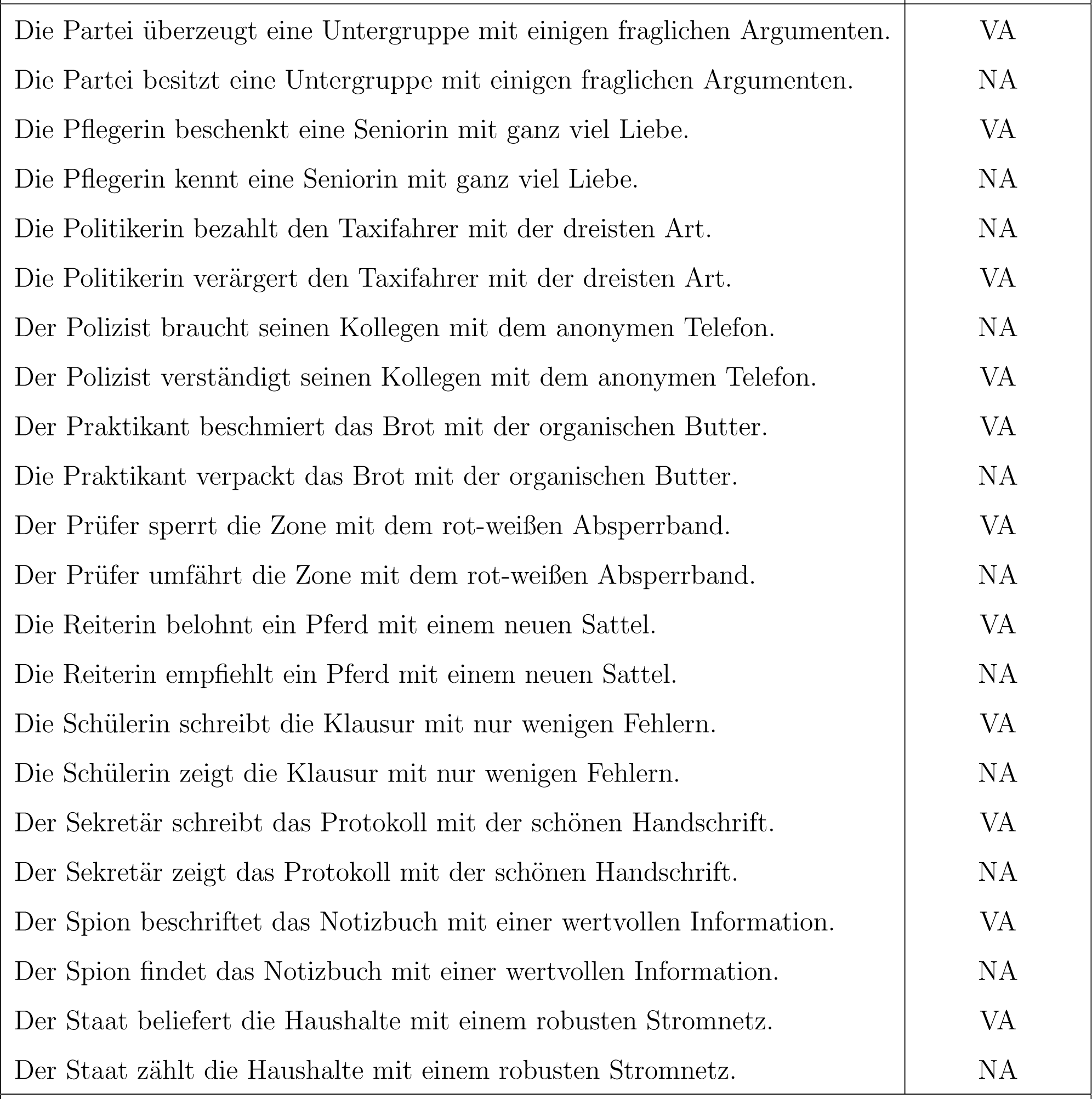
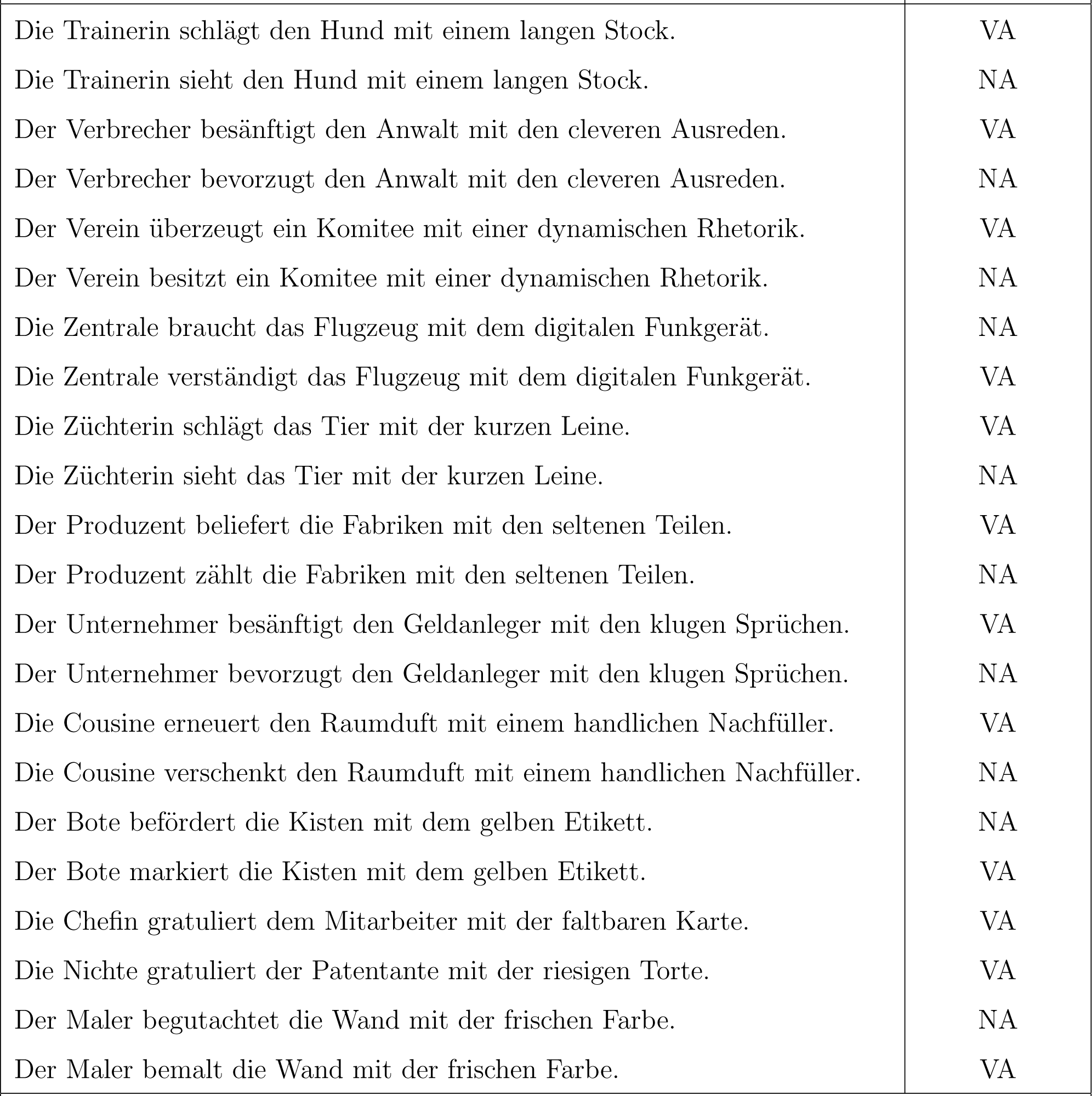
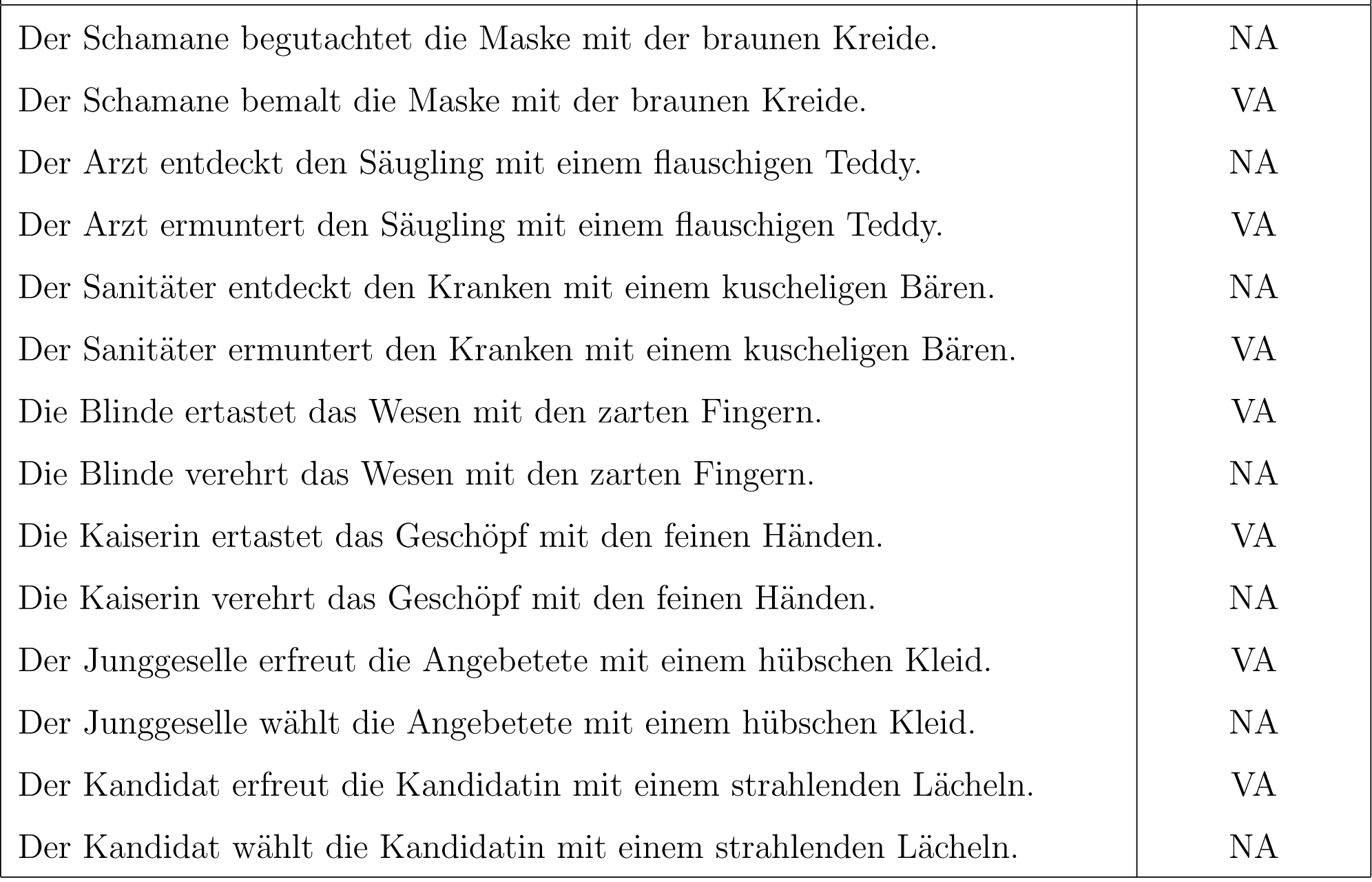
Stimulus Material - Verb condition

### Stimulus Material - Role condition

**Table 3:**
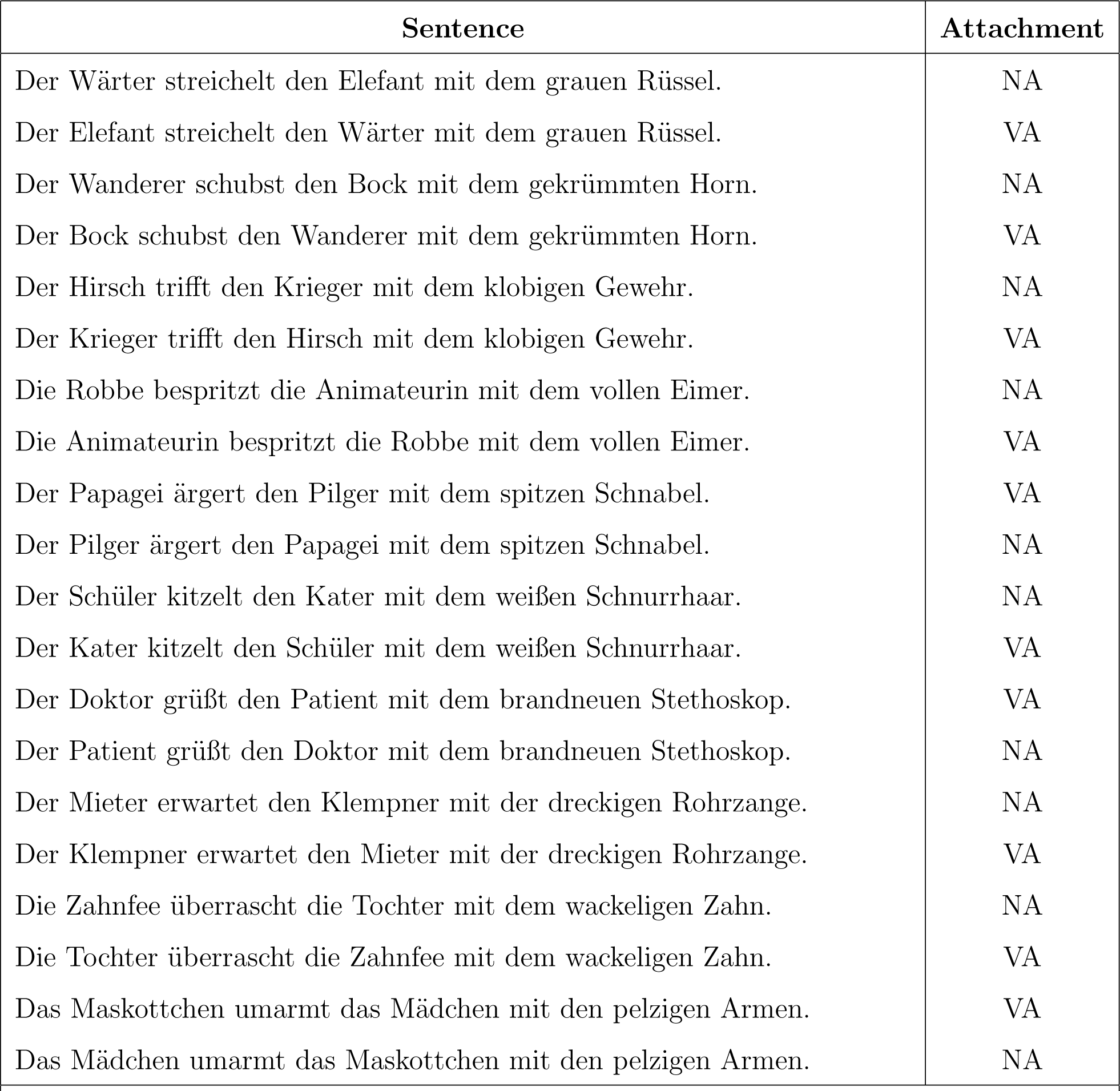
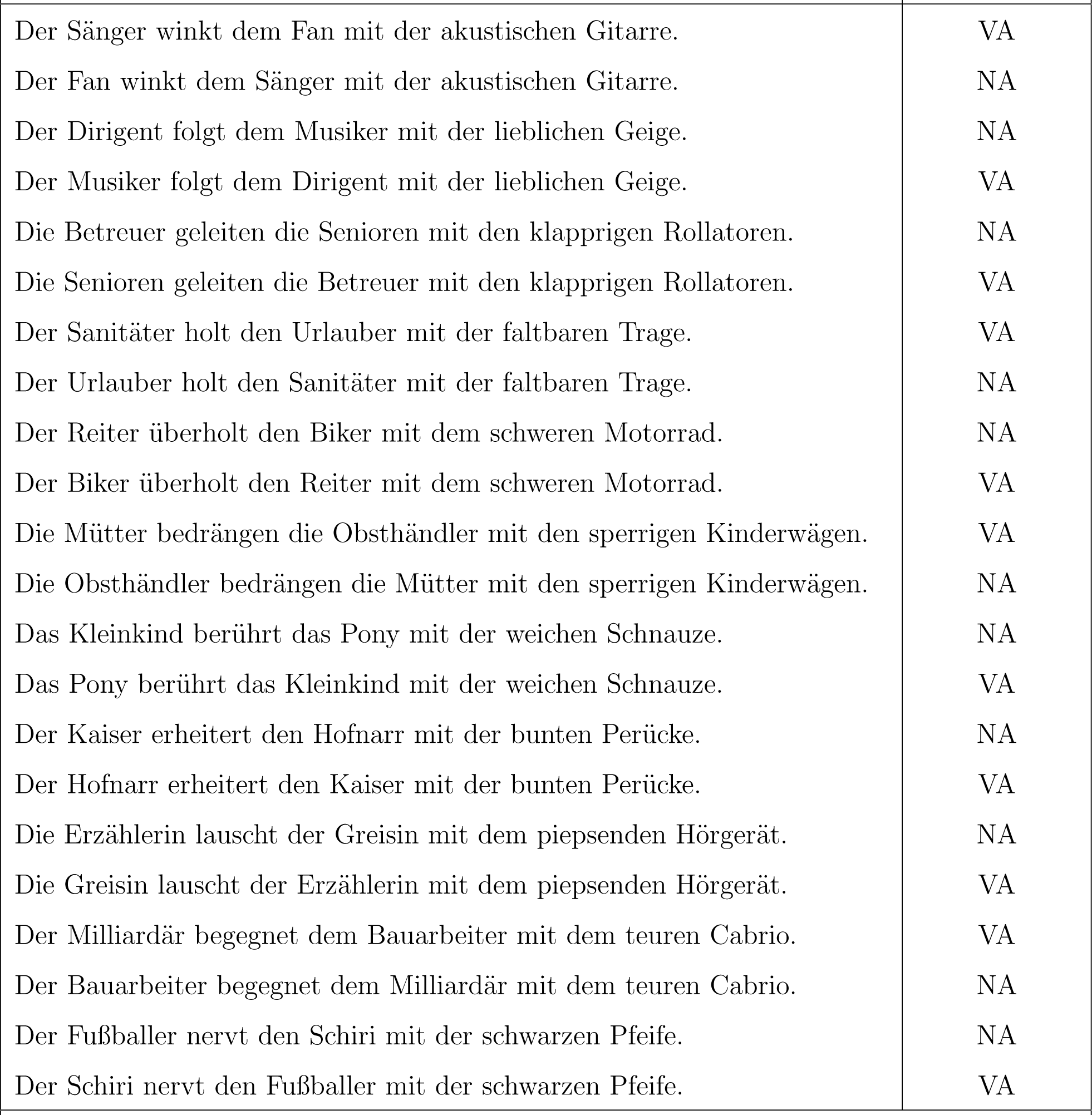
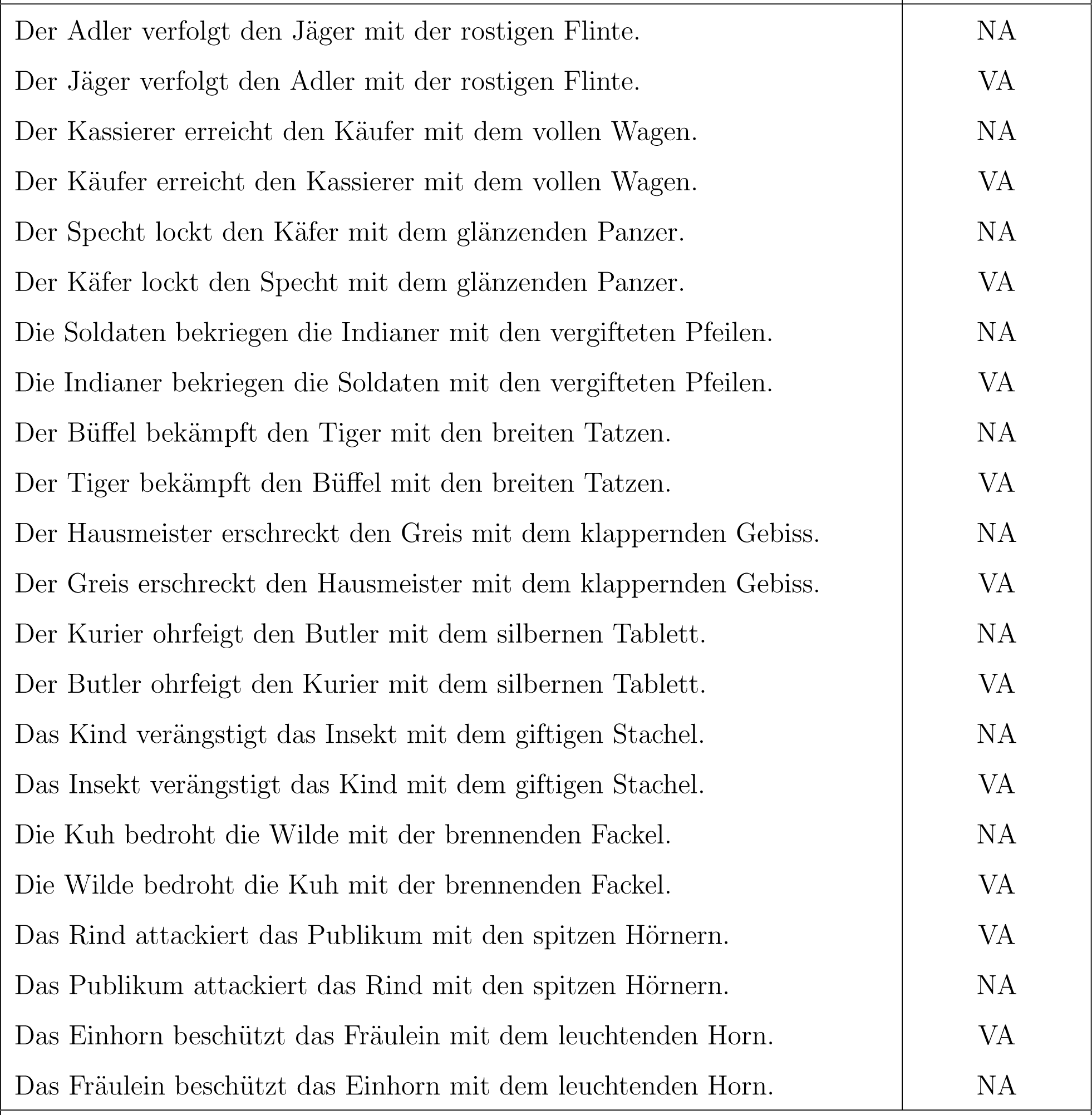
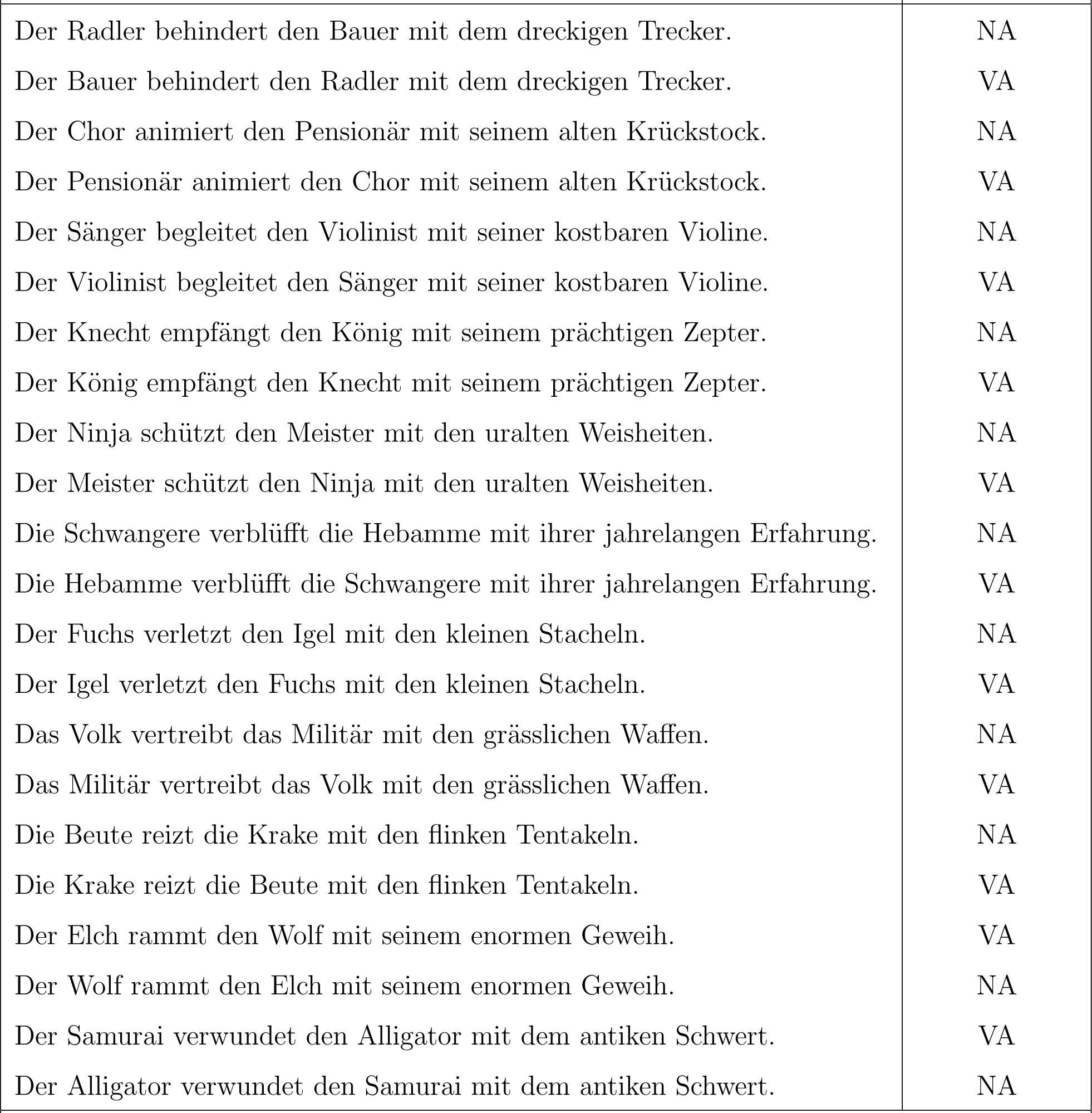
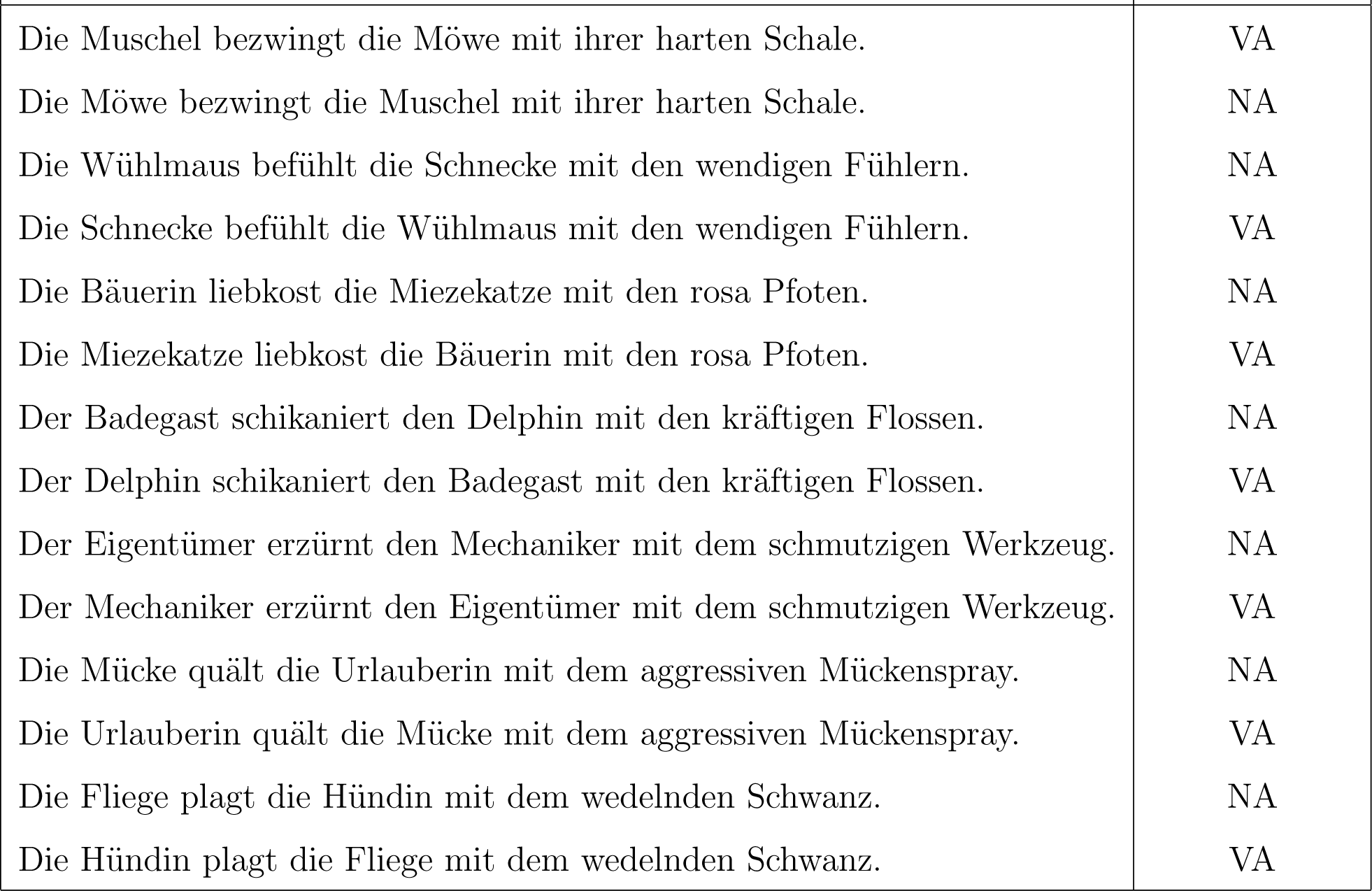
Stimulus Material - Role condition

## Footnotes

1 https://github.com/facebookresearch/fastText/blob/master/pretrained-vectors.md

2 Assuming that the thematic role is explicitly expressed and does not result from coercion

## References

1. Carsten Allefeld, Kai Görgen, and John Dylan Haynes. Valid population inference for information-based imaging: From the second-level t-test to prevalence inference. NeuroImage, 141:378–392, 2016. ISSN 10959572. doi: 10.1016/j.neuroimage.2016.07.040. URL http://dx.doi.org/10.1016/j.neuroimage.2016.07.040.

2. Gerry Altmann and Mark Steedman. Interaction with context during human sentence processing. Cognition, 30(3):191–238, 1988. ISSN 00100277. doi: 10.1016/0010-0277(88)90020-0.

3. D Bates, M Maechler, B Bolker, S Walker, R Haubo Bojesen Christensen, and Others. lme4: Linear mixed-effects models using Eigen and S4. R package version 1.1–7. 2014, 2015.

4. K Bock. Syntactic persistence in language processing. Cognitive Psychology, 18:355–387, 1986.

5. Piotr Bojanowski, Edouard Grave, Armand Joulin, and Tomas Mikolov. Enriching Word Vectors with Subword Information. 5:135–146, 2016. URL http://arxiv.org/abs/1607.04606.

6. Megan A Boudewyn, Megan Zirnstein, Tamara Y Swaab, and Matthew J Traxler. Priming prepositional phrase attachment: evidence from eye-tracking and event-related potentials. Quarterly journal of experimental psychology, 67(3):424–54, 2014. ISSN 1747-0226. doi: 10.1080/17470218.2013.815237. URL http://www.ncbi.nlm.nih.gov/pubmed/23859219.

7. Sabine Brants, Stefanie Dipper, Peter Eisenberg, Silvia Hansen-Schirra, Esther König, Wolfgang Lezius, Christian Rohrer, George Smith, and Hans Uszkoreit. TIGER: Linguistic Interpretation of a German Corpus. Research on Language and Computation, 2(4):597–620, 2004. ISSN 1572-8706. doi: 10.1007/s11168-004-7431-3. URL http://dx.doi.org/10.1007/s11168-004-7431-3.

8. Carsten Schmitz. LimeSurvey: An Open Source survey tool, 2012. URL http://www.limesurvey.org.

9. Radoslaw Martin Cichy, Dimitrios Pantazis, and Aude Oliva. Resolving human object recognition in space and time. Nature Neuroscience, 17(3): 455–462, 2014. ISSN 10976256. doi: 10.1038/nn.3635.

10. Cas W Coopmans, Helen De Hoop, Karthikeya Kaushik, Peter Hagoort, and Andrea E Martin. Structure-(in)dependent Interpretation of Phrases in Humans and LSTMs. Proceedings of the Society for Computation in Linguistics, 4(58), 2021.

11. Anders M Dale, Bruce Fischl, and Martin I Sereno. Cortical surface-based analysis: I. Segmentation and surface reconstruction. Neuroimage, 9(2): 179–194, 1999.

12. Tyler Davis, Karen F. LaRocque, Jeanette A. Mumford, Kenneth A. Norman, Anthony D. Wagner, and Russell A. Poldrack. What do differences between multi-voxel and univariate analysis mean? How subject-, voxel-, and trial-level variance impact fMRI analysis. NeuroImage, 97:271–283, 2014. ISSN 10959572. doi: 10.1016/j.neuroimage.2014.04.037. URL http://dx.doi.org/10.1016/j.neuroimage.2014.04.037.

13. Fernanda Ferreira and Matthew W. Lowder. Prediction, Information Structure, and Good-Enough Language Processing, volume 65. Elsevier Ltd, 2016. ISBN 9780128047903. doi: 10.1016/bs.plm.2016.04.002. URL http://dx.doi.org/10.1016/bs.plm.2016.04.002.

14. Fernanda Ferreira and Nikole D. Patson. The ‘Good Enough’ Approach to Language Comprehension. Language and Linguistics Compass, 1 (1-2):71–83, 2007. ISSN 1749-818X. doi: 10.1111/j.1749-818X.2007.00007.x. URL http://dx.doi.org/10.1111/j.1749-818X.2007.00007.x{%}5Cnhttp://doi.wiley.com/10.1111/j.1749-818X.2007.00007.x.

15. Hartmut Fitz and Franklin Chang. Sentence-level ERP effects as error propagation: A neurocomputational model. Cognitive Psychology, pages 1–46, 2018. doi: 10.31234/OSF.IO/FRX2W.

16. Stefan L. Frank and Rens Bod. Insensitivity of the human sentence-processing system to hierarchical structure. Psychological Science, 22(6): 829–834, 2011. ISSN 09567976. doi: 10.1177/0956797611409589.

17. Stefan L. Frank, Rens Bod, and Morten H. Christiansen. How hierarchical is language use? Proceedings of the Royal Society B: Biological Sciences, 279(1747):4522–4531, 2012. ISSN 14712954. doi: 10.1098/rspb.2012.1741.

18. Lyn Frazier and Keith Rayner. Making and correcting errors during sentence comprehension: Eye movements in the analysis of structurally ambiguous sentences. Cognitive psychology, 14(2):178–210, 1982.

19. Edouard Grave, Piotr Bojanowski, Prakhar Gupta, Armand Joulin, and Tomas Mikolov. Learning word vectors for 157 languages. In Proceedings of the International Conference on Language Resources and Evaluation (LREC 2018), 2018.

20. Tijl Grootswagers, Susan G. Wardle, and Thomas A. Carlson. Decoding Dynamic Brain Patterns from Evoked Responses: A Tutorial on Multivariate Pattern Analysis Applied to Time Series Neuroimaging Data. Journal of cognitive neuroscience, 29(4):677–697, 2017. ISSN 09528229. doi: 10.1162/jocn.

21. Matthias Guggenmos, Philipp Sterzer, and Radoslaw Martin Cichy. Multivariate pattern analysis for MEG: A comparison of dissimilarity measures. NeuroImage, 173(August 2017):434–447, 2018. ISSN 10959572. doi: 10.1016/j.neuroimage.2018.02.044. URL https://doi.org/10.1016/j.neuroimage.2018.02.044.

22. Peter Hagoort. The core and beyond in the language-ready brain. Neuroscience and Biobehavioral Reviews, 81:194–204, 2017. ISSN 18737528. doi: 10.1016/j.neubiorev.2017.01.048. URL https://doi.org/10.1016/j.neubiorev.2017.01.048.

23. Peter A. Hornby. Surface structure and presupposition. Journal of Verbal Learning and Verbal behavior, 13(5):530–538, 1974. ISSN 00225371. doi: 10.1016/S0022-5371(74)80005-8.

24. Ray Jackendoff. Pŕecis of Foundations of Language: Behavioral and Brain Sciences, 26(2003):651–707, 2003. ISSN 0140-525X. doi: 10.1017/S0140525X03000153.

25. T. Florian Jaeger. Categorical data analysis: Away from ANOVAs (transformation or not) and towards logit mixed models. Journal of Memory and Language, 59(4):434–446, 2008. ISSN 0749596X. doi: 10.1016/j.jml.2007.11.007. URL http://dx.doi.org/10.1016/j.jml.2007.11.007.

26. Koji Jimura and Russell A. Poldrack. Analyses of regional-average activation and multivoxel pattern information tell complementary stories. Neuropsychologia, 50(4):544–552, 2012. ISSN 00283932. doi: 10.1016/j.neuropsychologia.2011.11.007. URL http://dx.doi.org/10.1016/j.neuropsychologia.2011.11.007.

27. Nikolaus Kriegeskorte, Marieke Mur, and Peter A Bandettini. Representational similarity analysis-connecting the branches of systems neuroscience. Frontiers in systems neuroscience, 2:4, 2008.

28. Eric Maris and Robert Oostenveld. Nonparametric statistical testing of EEG- and MEG-data. Journal of Neuroscience Methods, 164(1):177–190, 2007. ISSN 01650270. doi: 10.1016/j.jneumeth.2007.03.024.

29. James L McClelland and Alan H Kawamoto. Mechanisms of sentence processing: Assigning roles to constituents. In Parallel distributed processing: explorations in the microstructure of cognition, vol. 2: psychological and biological models, pages 272–325. 1986.

30. Tom M Mitchell. Machine learning, volume 45. 1997.

31. Mohamed Taha Mohamed and Charles Jr. Clifton. Processing temporary syntactic ambiguity: The effect of contextual bias. Quarterly journal of experimental psychology (2006), 64(9):1797–1820, 2011. ISSN 1747-0218. doi: 10.1080/17470218.2011.582127.

32. Valeria Mongelli. The role of neural feedback in language unification: How awareness affects combinatorial processing. PhD thesis, Radboud University Nijmegen Nijmegen, 2020.

33. Marieke Mur, Peter A. Bandettini, and Nikolaus Kriegeskorte. Revealing representational content with pattern-information fMRI - An introductory guide. Social Cognitive and Affective Neuroscience, 4(1):101–109, 2009. ISSN 17495016. doi: 10.1093/scan/nsn044.

34. M. Nelson, I El Karoui, K Giber, X Yang, L Cohen, H Koopman, S Cash, L Naccache, J Hale, C Pallier, S Dehaene, Cognitive Neuroimaging Unit, and Los Angeles. Neurophysiological dynamics of phrase-structure building during sentence processing. pages 1–21, 2017. ISSN 0027-8424. doi: 10.1073/pnas.1701590114.

35. Guido Nolte. The magnetic lead field theorem in the quasi-static approximation and its use for magnetoenchephalography forward calculation in realistic volume conductors. Physics in Medicine and Biology, 48(22):3637– 3652, 2003. ISSN 00319155. doi: 10.1088/0031-9155/48/22/002.

36. Kenneth A. Norman, Sean M. Polyn, Greg J. Detre, and James V. Haxby. Beyond mind-reading: multi-voxel pattern analysis of fMRI data. Trends in Cognitive Sciences, 10(9):424–430, 2006. ISSN 13646613. doi: 10.1016/j.tics.2006.07.005.

37. Kayoko Okada, Feng Rong, Jon Venezia, William Matchin, I. Hui Hsieh, Kourosh Saberi, John T. Serences, and Gregory Hickok. Hierarchical organization of human auditory cortex: Evidence from acoustic invariance in the response to intelligible speech. Cerebral Cortex, 20(10):2486–2495, 2010. ISSN 10473211. doi: 10.1093/cercor/bhp318.

38. Robert Oostenveld, Pascal Fries, Eric Maris, and Jan Mathijs Schoffelen. FieldTrip: Open source software for advanced analysis of MEG, EEG, and invasive electrophysiological data. Computational Intelligence and Neuroscience, 2011:1–9, 2011. ISSN 16875273. doi: 10.1155/2011/156869.

39. Christophe Pallier, Anne-Dominique Devauchelle, and Stanislas Dehaene. Cortical representation of the constituent structure of sentences. Proceedings of the National Academy of Sciences of the United States of America, 108(6):2522–7, feb 2011. ISSN 1091-6490. doi: 10.1073/pnas.1018711108. URL http://www.pnas.org/cgi/content/long/108/6/2522.

40. Friedemann Pulvermüller, Yury Shtyrov, and Olaf Hauk. Understanding in an instant: Neurophysiological evidence for mechanistic language circuits in the brain. Brain and Language, 110(2):81–94, 2009. ISSN 0093934X. doi: 10.1016/j.bandl.2008.12.001. URL http://dx.doi.org/10.1016/j.bandl.2008.12.001.

41. Milena Rabovsky, Steven S Hansen, and James L McClelland. Modelling the N400 brain potential as change in a probabilistic representation of meaning. Nature Human Behaviour, 2(9):693–705, 2018. ISSN 23973374. doi: 10.1038/s41562-018-0406-4. URL http://dx.doi.org/10.1038/s41562-018-0406-4.

42. Rajeev D.S. Raizada, Feng Ming Tsao, Huei Mei Liu, and Patricia K. Kuhl. Quantifying the adequacy of neural representations for a cross-language phonetic discrimination task: Prediction of individual differences. Cerebral Cortex, 20(1):1–12, 2010. ISSN 10473211. doi: 10.1093/cercor/bhp076.

43. Keith Rayner, Marcia Carlson, and Lyn Frazier. The Interaction of Syntax and Semantics During Sentence Processing: Eye Movements in the Analysis of Semantically Biased Sentences. Journal of Verbal Learning and Verbal Behavior, 22(3):358–374, 1983.

44. Lilia Rissman and Asifa Majid. Thematic roles : Core knowledge or linguistic construct? Psychonomic Bulletin & Review, 2019.

45. Irina Simanova, Marcel A J van Gerven, Robert Oostenveld, and Peter Hagoort. Predicting the Semantic Category of Internally Generated Words from Neuromagnetic Recordings. Journal of Cognitive Neuroscience, 27(1):35–45, 2015. doi: 10.1162/jocn_a_00690. URL https://doi.org/10.1162/jocn{_}a{_}00690.

46. Michael Spivey-Knowlton and Julie C Sedivy. Resolving attachment ambiguities with multiple constraints. Cognition, 55(3):227–267, 1995. ISSN 00100277. doi: 10.1016/0010-0277(94)00647-4.

47. Arjen Stolk, Ana Todorovic, Jan Mathijs Schoffelen, and Robert Oostenveld. Online and offline tools for head movement compensation in MEG. NeuroImage, 68:39–48, 2013. ISSN 10538119. doi: 10.1016/j.neuroimage.2012.11.047. URL http://dx.doi.org/10.1016/j.neuroimage.2012.11.047.

48. Roman Taraban and James L McClelland. Constituent Attachment and Thematic Role Assignment in Sentence Processing: Influences of content-based expectations. Journal of Memory and Language, 27:597–632, 1988.

49. Matthew J. Traxler. Trends in syntactic parsing: Anticipation, Bayesian estimation, and good-enough parsing. Trends in Cognitive Sciences, 18 (11):605–611, 2014. ISSN 1879307X. doi: 10.1016/j.tics.2014.08.001. URL http://dx.doi.org/10.1016/j.tics.2014.08.001.

50. Matthew J. Traxler and Kristen M. Tooley. Lexical mediation and context effects in sentence processing. Brain Research, 1146(1):59–74, 2007. ISSN 00068993. doi: 10.1016/j.brainres.2006.10.010.

51. Lorraine K. Tyler, Teresa P.L. Cheung, Barry J. Devereux, and Alex Clarke. Syntactic computations in the language network: Characterizing dynamic network properties using representational similarity analysis. Frontiers in Psychology, 4(MAY):1–19, 2013. ISSN 16641078. doi: 10.3389/fpsyg.2013.00271.

52. Daniëlle Van Den Brink and Peter Hagoort. The influence of semantic and syntactic context constraints on lexical selection and integration in spoken-word comprehension as revealed by ERPS. Journal of Cognitive Neuroscience, 16(6):1068–1084, 2004. ISSN 0898929X. doi: 10.1162/0898929041502670.

53. Mats WJ van Es, Tom R Marshall, Eelke Spaak, Ole Jensen, and Jan-Mathijs Schoffelen. Phasic modulation of visual representations during sustained attention. European Journal of Neuroscience, 2020.

54. M Van Gerven, A Bahramisharif, J Farquhar, and T Heskes. Donders Machine Learning Toolbox (DMLT). https://github.com/distrep/DMLT, version, 26(06):2013, 2013.

55. B. D. Van Veen, W van Drongelen, M Yuchtman, and A Suzuki. Localization of brain electrical activity via linearly constrained minimum variance spatial filtering. IEEE Transactions on Biomedical Engineering, 44(9): 867–880, 1997. ISSN 0018-9294. doi: 10.1109/10.623056.

56. Gaël Varoquaux. Cross-validation failure: Small sample sizes lead to large error bars. NeuroImage, (April):1–10, 2017. ISSN 10959572. doi: 10.1016/j.neuroimage.2017.06.061.

57. Diego Vidaurre, Nicholas E. Myers, Mark Stokes, Anna C. Nobre, and Mark W. Woolrich. Temporally Unconstrained Decoding Reveals Consistent but Time-Varying Stages of Stimulus Processing. Cerebral Cortex, 29(2):863–874, 2019. ISSN 14602199. doi: 10.1093/cercor/bhy290.

58. Michael J. Wolff, Janina Jochim, Elkan G. Akyürek, and Mark G. Stokes. Dynamic hidden states underlying working-memory-guided behavior. Nature Neuroscience, 20(6):864–871, 2017. ISSN 15461726. doi: 10.1038/nn.4546.

59. Atsushi Yokoi and Jörn Diedrichsen. Neural Organization of Hierarchical Motor Sequence Representations in the Human Neocortex. Neuron, 103(6): 1178–1190.e7, 2019. ISSN 10974199. doi: 10.1016/j.neuron.2019.06.017.

60. Jayden Ziegler, Giulia Bencini, Adele Goldberg, and Jesse Snedeker. How abstract is syntax? Evidence from structural priming. Cognition, 193(December 2018):104045, 2019. ISSN 18737838. doi: 10.1016/j.cognition.2019.104045. URL https://doi.org/10.1016/j.cognition.2019.104045.

